# Metal-independent ribonucleotide reduction powered by a DOPA radical in Mycoplasma pathogens

**DOI:** 10.1101/348268

**Authors:** Vivek Srinivas, Hugo Lebrette, Daniel Lundin, Yuri Kutin, Margareta Sahlin, Michael Lerche, Jürgen Eirich, Rui M. M. Branca, Nicholas Cox, Britt-Marie Sjöberg, Martin Högbom

## Abstract

Ribonucleotide reductase (RNR) catalyzes the only known de-novo pathway for production of all four deoxyribonucleotides required for DNA synthesis. In aerobic RNRs, a di-nuclear metal site is viewed as an absolute requirement for generating and stabilizing an essential catalytic radical. Here we describe a new group of RNRs found in Mollicutes, including Mycoplasma pathogens, that possesses a metal-independent stable radical residing on a modified tyrosyl residue. Structural, biochemical and spectroscopic characterization reveal a stable DOPA radical species that directly supports ribonucleotide reduction *in vitro* and *in vivo*.

The cofactor synthesis and radical generation processes are fundamentally different from established RNRs and require the flavoprotein NrdI. Several of the pathogens encoding this RNR variant are involved in diseases of the urinary tract and genitalia. Conceivably, this remarkable RNR variant provides an advantage under metal starvation induced by the immune system. We propose that the new RNR subclass is denoted class Ie.

## Main text

Ribonucleotide reductase (RNR) catalyzes the only known de-novo pathway for production of all four deoxyribonucleotides required for DNA synthesis (*1, 2*). It is essential for all organisms with DNA as genetic material and a current drug target (*3, 4*). In 1968 it was discovered that non-heme iron is required for function in aerobic, class I, RNR found in all eukaryotes and many bacteria (*5, 6*). Since then, through half a century of intense studies, a di-nuclear metal site has been viewed as an absolute requirement for generating and stabilizing a catalytic radical, essential for RNR activity (*7*).

In all domains of life ribonucleotides are reduced to their corresponding 2’-deoxyribonucleotides in a radical-catalyzed reaction. Three RNR classes have been discovered, all requiring transition metals for function (*2*): Class III is strictly anaerobic and use a 4Fe-4S cluster for radical generation. Class II is indifferent to oxygen and utilizes an adenosyl cobalamin cofactor. In all hitherto studied class I RNRs, the catalytic radical is generated and stabilized by a dinuclear metal site in protein R2 in an oxygen dependent reaction, and then reversibly shuttled to protein R1 where ribonucleotide reduction occurs (8-10). The dinuclear metal site is coordinated by four carboxylate residues and two histidines. Depending on subclass, the active form of the metal site can be either di-iron (class Ia), di-manganese (class Ib), or heterodinuclear Mn/Fe (class Ic) (11, 12). Classes Ia and Ib generate a stable tyrosyl radical that is magnetically coupled to the metal site (*13, 14*). Class Ic instead forms a radical-equivalent Mn^IV^/Fe^III^ high-valent oxidation state of the metal site (*15, 16*). A class Id, containing a Mn^IV^/Mn^III^ cofactor, was also recently proposed (17-19). The dinuclear metal sites in classes Ia and Ic perform direct oxygen activation while class Ib requires an additional flavoprotein, NrdI, to generate superoxide used to oxidize the dimanganese site, ultimately resulting in tyrosyl radical generation (*20-22*).

Here we describe a new group of RNR proteins in Mollicutes, including Mycoplasma pathogens, that possesses a metal-independent stable radical residing on a modified tyrosyl residue. Structural, biochemical and spectroscopic characterization reveal an unprecedented and remarkably stable DOPA radical species that directly supports ribonucleotide reduction *in vitro* and *in vivo.* We propose that the new RNR subclass is denoted class Ie.

### A new RNR subclass able to rescue an Escherichia coli strain lacking aerobic RNR

Sequence analysis revealed a group of class I RNR operons, present in common human pathogens e.g. *Mycoplasma genitalium*, *Mycoplasma pneumoniae*, and *Streptococcus pyogenes*. Analogous to standard class Ib RNRs, the operons contain the genes *nrdE*, *nrdF* and *nrdI*, coding for proteins R1, R2 and NrdI, respectively. Phylogenetically, the group forms a clade derived from class Ib proteins (Fig S1). Strikingly, unlike any previously identified *nrdE/F* genes, the R2 proteins encoded by this group retain only 3 of the 6 metal binding residues that are otherwise completely conserved, and individually essential, regardless of R2 subclass. Two variants are observed in which 3 of the 4 carboxylate metal ligands are either substituted for valine, proline and lysine (VPK variant) or for glutamine, serine and lysine (QSK variant) (Fig 1A). In both cases, the substitutions render the site formally charge neutral, a drastic change from the −4 net charge of the metal binding site in other R2 proteins. These substitutions appear to exclude a metal site and a radical generation mechanism even remotely similar to any thus far studied ribonucleotide reductase. Apart from the metal binding site, sequence and structural analysis of the R1, R2 and NrdI proteins show that the residues normally required for activity, including the R1 active site and the radical transfer path between the R1 and R2 subunits are conserved. As common for class I RNRs, several genomes encoding the QSK or VPK variant also harbor other RNRs (Figure 1B). However, the QSK/VPK variants also often represent the only aerotolerant RNR found in the genome, for example in *Mycoplasma genitalium* and *Mycoplasma pneumoniae* (VPK) and *Gardnerella vaginalis* (QSK).

We investigated if a VPK variant operon could rescue an *E. coli* strain lacking aerobic RNR (Δ*nrdAB*, Δ*nrdEF*) (*23*), thus otherwise unable to grow in the presence of oxygen. A tunable arabinose-induced pBAD plasmid containing the *nrdFIE* operon from *Mesoplasma florum* was constructed and transformed into the (Δ*nrdAB*, Δ*nrdEF*) strain. Cultures grown under anaerobic conditions were subsequently exposed to oxygen. As shown in figure 1C, the (Δ*nrdAB*, Δ*nrdEF*) strain did not recover growth under aerobic conditions while the strain transformed with the *MfnrdFIE* containing plasmid recovered growth when expression of the operon was induced by arabinose, but not when suppressed by glucose. We conclude that the operon can rescue this knockout strain and provide competence to grow under aerobic conditions. This result is consistent with our previous observations of the *Streptococcus pyogenes nrdFIE* operon (*24*).

### *MfNrdI* activates *Mf*R2 in an O_2_ dependent reaction

We proceeded to quantify the *in vitro* activity of the enzyme. We were unable to obtain *in vitro* ribonucleotide reductase activity using the *Mf*R2 protein expressed separately in *E. coli*. However, purification of *Mf*R2 after co-expression of the entire *MfnrdFIE* operon under aerobic conditions rendered a deep-blue protein that exhibited RNR activity together with protein *Mf*R1 (Fig 1D). The color and *in vitro* RNR activity was also observed when *Mf*R2 was co-expressed with only *MfnrdI* under aerobic conditions while co-expression under anaerobic conditions rendered an inactive and colorless protein R2. Under the current assay conditions, we calculate a turnover number of 0.18 s^−1^ or > 300 for the duration of the assay. Once the R2 protein is activated, *Mf*NrdI is thus not required for multi turnover activity *in vitro*. The specific activity was determined to 275 ± 7 nmo1×min^−1^×mg^−1^, on par with typical class I RNRs (*18, 22*). Incubation of the active R2 protein with hydroxyurea, a radical-quenching RNR inhibitor, rendered a colorless and inactive protein. The activity of the quenched protein could be partially restored if MfNrdI was added to the inactivated *Mf*R2 and subjected to reduction-oxidation cycles using dithionite and oxygen-containing buffer (Fig 1D).

Together, the data show that despite the substitutions of the metal binding residues, *Mf*R2 and *Mf*R1 constitute an active RNR system, but only after protein R2 has undergone an NrdI and O_2_ dependent activation step. This is principally similar to class Ib RNRs. Moreover, Small Angle X-ray Scattering (SAXS) measurements showed that *Mf*R2 and *Mf*NrdI forms a well-defined 2:2 complex with the same interaction geometry as previously observed in standard class Ib RNR proteins (Fig. S2) (*21, 25*). Remarkably, however, for class Ib RNR, the role of NrdI is to provide an oxidant for the di-manganese metal site that subsequently generates the catalytic tyrosyl radical. A mechanism that appears implausible, given the substitution of the metal binding residues in *Mf*R2.

**Fig 1.**
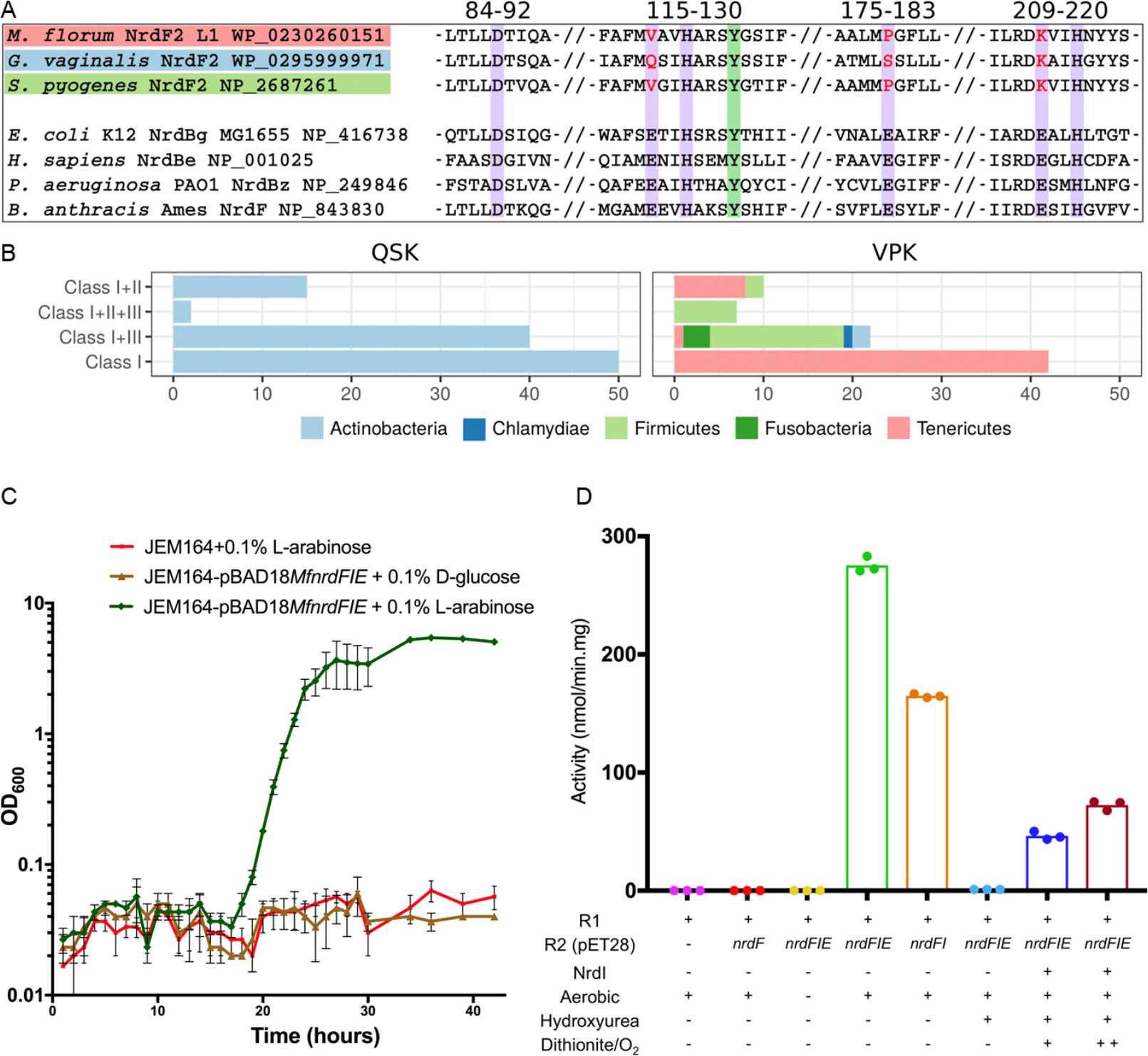
Features and activity of the new RNR subgroup. **A)** Sequence alignment of the new R2 protein groups to a number of standard, di-metal containing, R2 proteins. Purple background indicates the 6 normally essential metal-binding residues, only 3 of which are conserved. The normally radical harboring tyrosine residue is shown with a green background. **B)** Taxonomic distribution of NrdF2. QSK/VPK encoding organisms and their collected RNR class repertoire. The QSK clusters are found only in Actinobacteria, whereas the VPK clusters are also found in Firmicutes, Tenericutes, Chlamydiae and Fusobacteria. **C)** Expression of the *MfnrdFIE* operon induced by addition of 0.1% v/v L-arabinose (green) rescued the double knockout strain while when gene expression was suppressed with 0.1% v/v D-glucose (brown) the strain failed to recover, as did the strain lacking the vector (red). **D)** HPLC based *in vitro* RNR activity assays show no activity for R2 protein expressed separately (red), while co-expression with *MfnrdI* and *MfnrdE* (green) or *MfnrdI* (orange) produced an active R2 protein. Anaerobic co-expression (yellow) or incubation of the active R2 with hydroxyurea (light blue) abolishes the activity. Partial activity could be regenerated by the addition of *Mf*NrdI and redox cycling with dithionite and oxygen, blue and maroon for one and two reduction-oxidation cycles respectively.

### The active R2 protein is metal-free but covalently modified

The crystal structure of *Mf*R2 in its active form (co-expressed with the entire RNR operon under aerobic conditions) was determined to 1.5 Å resolution. In addition, two crystal structures of the inactive form, either expressed alone aerobically or co-expressed with the entire operon under anaerobic conditions, were determined to 1.2 Å resolution (Table S1).

Overall, the three structures show the same fold and dimeric arrangement as standard metalbinding R2 proteins (Fig 2A).

**Fig 2.**
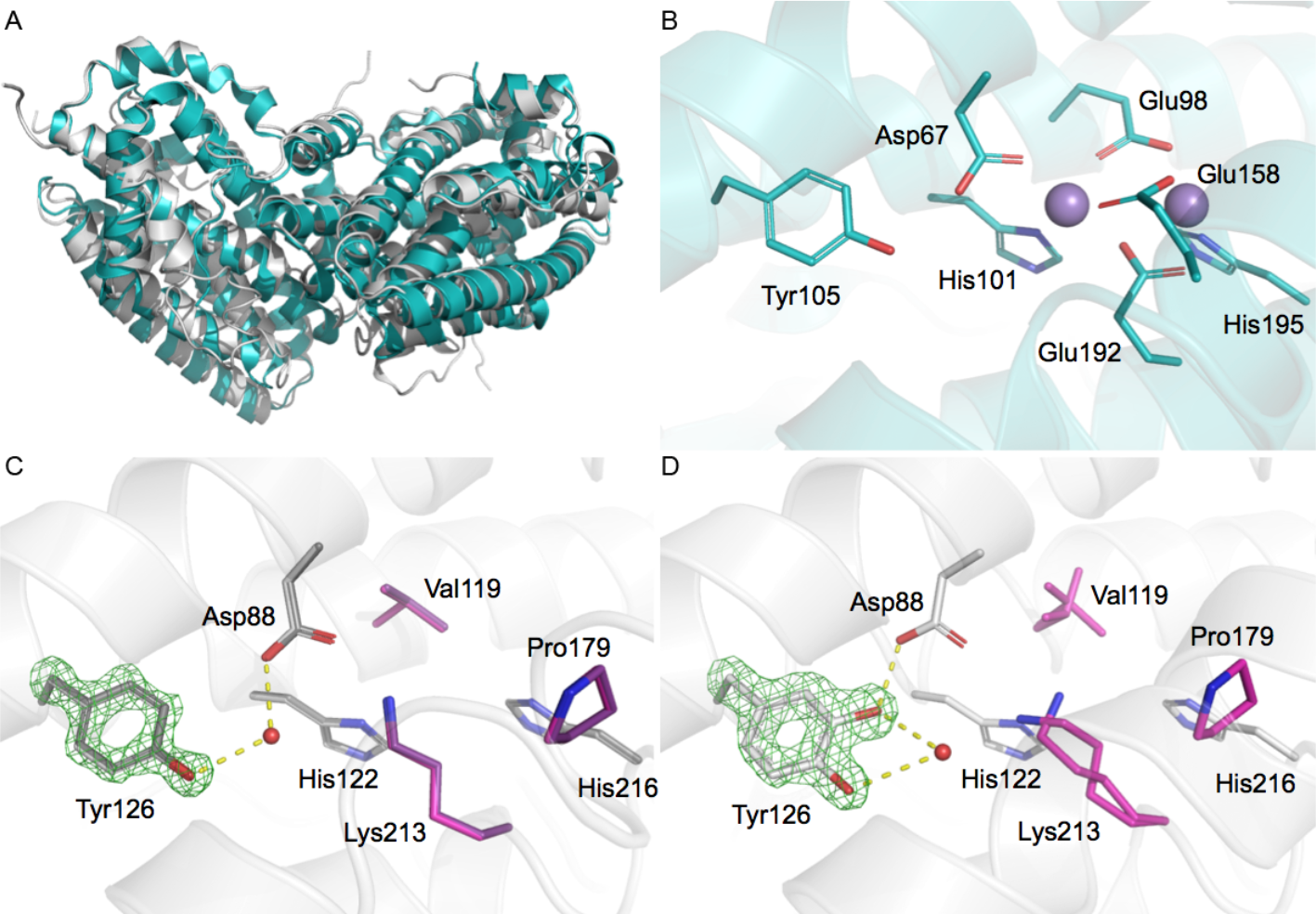
Crystal structures of inactive and active forms of *Mf*R2. **A)** Overall structure of the *Mesoplasma florum* VPK R2 protein (gray) compared to the standard class Ib R2 from *Escherichia coli* (PDB: 3n37) (cyan). **B)** Structure of the dinuclear metal site and conserved metal coordinating residues in standard class I RNR R2. **C)** Structure of inactive *Mf*R2 after expression without *Mf*Nrdl (gray) or with M/NrdI under anaerobic conditions (dark gray), these structures are identical within experimental error. The 3 residues substituting the normally conserved carboxylate metal-ligands in canonical class I R2s are shown in pink and purple respectively. **D)** Structure of the active *Mf*R2 after aerobic co-expression with NrdI and R1. Simulated annealing omit Fo-Fc electron density maps for the unmodified or modified Tyr126 is shown in green and contoured at 8σ. Carbons are in cyan, gray/pink for R2 from *E. coli* and *M. florum*, respectively. Oxygens and nitrogens are colored red and blue, respectively. Mn(II) ions are represented as purple spheres. H-bond interactions to the Tyr126 are indicated.

As expected from sequence, the site normally occupied by the di-nuclear metal cofactor is remarkably atypical with the substitution of three of the six otherwise conserved and individually essential metal coordinating residues, leaving only one carboxylate and two histidines (Fig. 2B,C). No electron density corresponding to a metal ion could be observed in the site in any of the structures. We also collected X-ray anomalous scattering data at a wavelength of 0.97 Å and calculated anomalous difference maps, a method which would detect even low occupancy metal binding. Still, no signal above noise could be observed in the vicinity of the site. To rule out that metal was lost during crystallization, we conducted Total-reflection X-Ray Fluorescence (TXRF) analysis on the active *Mf*R2 protein solution, quantitatively detecting all elements from aluminum to uranium (with the exception of Zr, Nb, Mo, Tc, Ru and Rh). Only trace amounts of transition metals could be detected in the active protein sample (e.g. mol/mol metal/protein: Mn 0.04%, Fe 0.35%, Co 0.00%, Ni 1.70%, Cu 0.99%, Zn 0.71%, (Fig. S3 and Table S2)). Cumulatively, the metal content corresponded to less than 0.04 per protein monomer with nickel being the dominating species, most likely due to that the purification involved a Ni-affinity step. TXRF analyses were also performed for the purified *Mf*R1 and *Mf*NrdI proteins in solution, again with only trace amounts of metals detected.

In the *Mf*R2 crystal structure, the canonical metal positions are occupied: 1) by a water molecule in a tetrahedral coordination, involving the conserved His216, with distances of 2.8 ± 0.1 Å, as expected for a hydrogen-bonded water, but very unlikely for a metal; 2) by the ε-amino group of Lys213, replacing the conserved metal-bridging glutamate present in all class I R2s. This lysine forms a hydrogen bond with Asp88, the only remaining carboxylate residue. Asp88 also interacts via a H-bonded water with Tyr126, corresponding to the tyrosine harboring the metal-coupled radical in standard class Ia and Ib R2 proteins (Fig. 2C).

Notably, in the active protein, the tyrosine appears covalently modified in the meta-position, a modification not observed in any of the two 1.2 Å crystal structures of the inactive *Mf*R2 proteins (Fig. 2C,D). Mass spectrometry analysis confirmed that the protein is covalently modified. The full-length active protein is 17 ± 2 Da heavier than the inactive *Mf*R2, 39804 ± 2 Da vs. 39787 ± 1.4 Da, respectively (Fig S4). Proteolytic cleavage and peptide analysis pinpoints the modification to + 15.995 ± 0.003 Da (monoisotopic mass, corresponding to one additional oxygen atom) at Tyr126 (Fig S5). Based on the X-ray crystallographic and mass spectrometry data, we thus conclude that Tyr126 is meta-hydroxylated in the active protein.

### Characterization of a novel stable DOPA radical species in *Mf*R2

The tyrosyl radical in canonical R2 proteins shows a sharp UV/vis absorbance peak at ≈ 410 nm. The active *Mf*R2 shows a peak with a similar profile though significantly blue-shifted to ≈ 383 nm with substantial additional structure at lower wavelengths (Fig. 3A). It has previously been demonstrated in simpler phenoxyl radical model systems that meta OH substitution leads to a blue shift of the radical marker band (*26*), suggesting that a radical is located on the modified tyrosine residue. No such absorbance was observed in the inactive *Mf*R2. Incubation of 60 μM protein with 52 mM hydroxyurea for 20 minutes led to the complete disappearance of the absorbance features and rendered a colorless protein. The rate constants for the decay of the absorbance at 348, 364 and 383 nm was identical, consistent with all absorbance features arising from a single radical species (Fig. 3A). In absence of radical quenchers the features are remarkably stable with no observable decay during 400 minutes at 25°C (Fig. S6).

**Fig 3.**
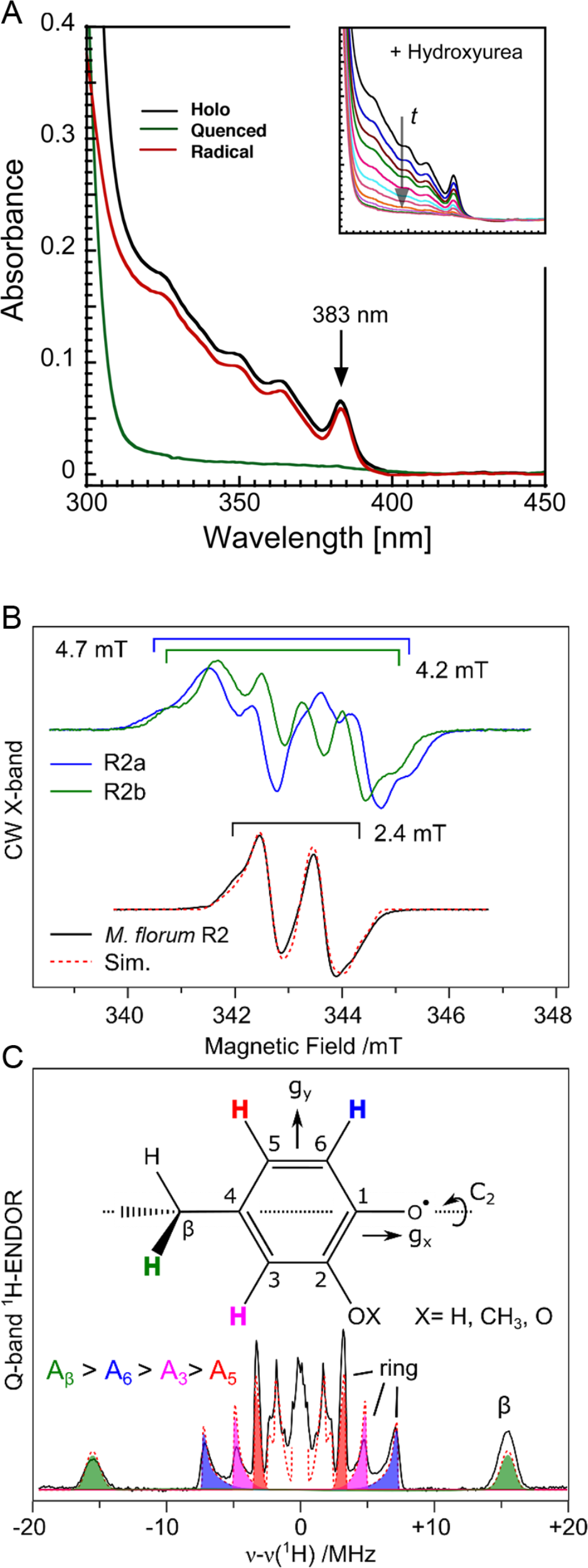
Spectroscopic characterization. **A)** The UV/vis spectrum of the active blue protein shows a peak at 383 nm and additional structure at lower wavelengths (black). Incubation with 52 mM hydroxyurea for 20 minutes removed all features from the spectrum except the protein-related absorbance peak at 280 nm (green and inset). The red trace represents the active minus quenched spectra. **B)** X-band EPR spectra of tyrosyl radicals observed in R2 proteins (*Escherichia coli* R2a and *Bacillus cereus* R2b reconstituted with Fe) compared to the *M. florum* R2 radical species reported here. **C)** Q-band ENDOR spectrum recorded at the low field edge of the EPR spectrum. The red dashed lines represent a simultaneous simulation of all datasets. The radical species is assigned to the structure shown.

We further used Electron Paramagnetic Resonance (EPR) spectroscopy to conclusively demonstrate that this new class of R2 proteins contains a radical. The identity of the radical can be determined from its spectral shape and width. These features stem from the coupling of the unpaired electron spin (S = ½) to hydrogen nuclei (protons, I = ½) in its vicinity via hyperfine interactions. Each interaction can also be measured separately using the complementary technique Electron Nuclear DOuble Resonance (ENDOR). From inspection of the EPR spectrum it can be readily seen that the width of the signal is too narrow to be assigned to an unmodified tyrosyl radical (Figure 3B,C, for a full description see SI text, Fig S7-S9 and tables S3-S5). Spectral simulations of the multifrequency EPR and ENDOR spectra using the spin Hamiltonian formalism reveal that the radical has four non-equivalent proton couplings with isotropic values of 28.7, 9.8 6.5 and 4.4 MHz. The largest coupling is assigned to a proton of the C4 alkyl substituent (Cβ). The remaining three couplings are assigned to protons of the conjugated ring. As no equivalent proton couplings are observed, the ring must be asymmetric, requiring the introduction of an additional ring substituent, consistent with the C2 substitution observed in the crystal structure. The overall decrease in coupling constants as compared to literature tyrosyl radicals suggests that the new substituent is an oxy group (O-X) (Fig. 3C) (*27, 28*), although it is noted that the unpaired spin density associated with the oxy groups of C1 and C2 is not evenly shared, as would be the case for an *o*-semiquinone(*29*). X cannot be conclusively determined from the EPR/ENDOR measurements but is fully consistent with a hydroxyl substituent or a strong hydrogen bond as indicated by crystallography and mass spectrometry (for further discussion see SI).

Unlike typical R2 proteins, the radical displays relaxation behavior corresponding to no (or weak) interaction with a metal (Fig. S10), consistent with metal analysis and crystallographic data. Instead the evaluated power of half saturation more closely coincides with that for a tyrosyl radical in solution generated by light illumination (*8, 30, 31*). Finally, quantification of the radical species by EPR shows that the active *Mf*R2 sample contains ≈ 52% of radical per protein monomer. As described previously, the cumulative amount of transition metals measured by TXRF for the same sample is less than 4% per protein. It is thus not possible that a metal ion is required for stabilization of the observed radical species.

Comprehensive structural, EPR, UV/vis, TXRF and mass spectrometry data thus support that the novel radical species is metal-independent, located on a meta-hydroxylated tyrosyl residue and represents an unprecedented intrinsic DOPA radical within an RNR system.

### The observed radical species supports ribonucleotide reduction

To investigate whether the observed radical species is catalytically competent, we utilized the mechanism-based inhibitor 2’-azido-2’deoxycytidine-5’-diphosphate (N_3_-CDP). It has previously been shown that incubation of RNR with N_3_-CDP under turnover conditions leads to the loss of the R2 radical, concomitant with the formation of a nitrogen-centered radical in the active site of protein R1 (*32-34*). Incubation of active *Mf*R2 with *Mf*R1 and N_3_-CDP led to the quantitative disappearance of the R2-centered radical species. Catalytic turnover with the substrate CDP or incubation of protein R2 with N_3_-CDP in the absence of protein R1 did not lead to quenching of the R2 radical (Fig. 4A). These results prove that the observed metal-independent radical species in protein*Mf*R2 initiates the catalytic radical chemistry in protein R1 in this new RNR subclass.

**Fig 4.**
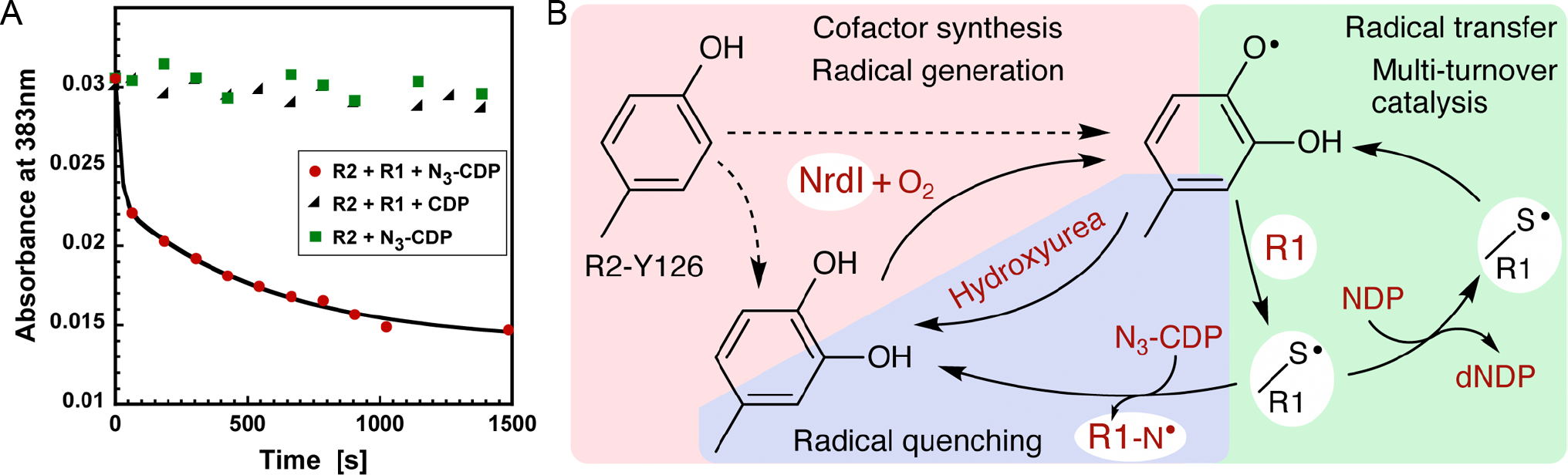
Catalytic competency of the radical and proposed mechanistic scheme in class le RNR. **A)** The radical signal is quenched in the presence of protein R1 and the mechanism-based inhibitor N_3_-CDP (red). Protein R2 with N_3_-CDP alone (green) or turnover conditions with protein R1 and CDP (black) does not quench the *Mf*R2 radical. The experiment shows that the observed radical can be reversibly transferred to the active site and support catalysis in protein R1. **B)** Proposed mechanistic steps in class Ie RNR. Dashed lines indicate alternative paths for the NrdI and O_2_-dependent posttranslational modification of Tyr126.

## Discussion

Here we identify a novel type of class I RNRs, found in human pathogens, which performs aerobic multiple-turnover ribonucleotide reduction using a metal-independent radical. We propose that this RNR group is denoted class Ie. Figure 4B summarizes our current understanding of the system. The radical-harboring cofactor is first post-translationally generated by hydroxylation of Tyr126 in an NrdI and O_2_ dependent process. It is presently unknown if this reaction also directly forms the radical species or if it is always generated in a subsequent NrdI-dependent step. Once the DOPA radical is formed in protein R2 it supports multi-turnover ribonucleotide reduction together with protein R1, presumably analogous to other class I RNR systems. It has previously been shown that a synthetically induced DOPA residue in the radical transfer path in *E. coli* RNR functions as a radical trap preventing RNR turnover, likely due to a too low reduction potential (*35*). The EPR characteristics of the stable radical in *Mf*R2 show that it is electronically more similar to a substituted tyrosyl radical than an *o*-semiquinone. It appears likely that this asymmetry is induced by the protein to tune the redox potential allowing reversible transfer to the R1 subunit.

If the radical is lost, activity can be restored in the covalently modified R2 protein by NrdI. Remarkably, NrdI achieves two different oxygen-dependent reactions in this system, tyrosine hydroxylation and DOPA radical generation. Flavoproteins are known to be able to produce two different O_2_-derived reactive oxygen species, hydrogen peroxide and superoxide. There is chemical precedent for hydrogen peroxide/hydroxyl radical tyrosine hydroxylation as well as DOPA radical generation by superoxide (*36-38*). Thus, an intriguing possibility is that both these oxidants are utilized at different stages of activation of the R2 protein.

Ribonucleotide reductase catalyzes the only known de-novo pathway for production of all four deoxyribonucleotides and was the first enzyme identified to utilize a protein-derived radical for catalysis (*31, 39*). Half a century ago it was established that aerobic RNR requires iron for function (*5*). Since then, two other classes of RNR have been discovered and while the radical generation systems differ, all types of RNR, in all domains of life, have been considered to have an absolute metal dependence in order to generate the essential catalytic radical. Hitherto studied class I RNRs requires oxygen and a dinuclear metal site, which depending on subclass is: di-iron, di-manganese, or heterodinuclear manganese/iron. Differences in metal requirement among RNRs, of which many organisms possess more than one, are believed to provide an advantage in environments where some metals are limiting. Metal starvation, including restricting access to both iron and manganese, is a central strategy in innate immunity to combat invading pathogens (*40*). It is tempting to speculate that class Ie RNR evolved in response to such extremely metal-restricted environments. The post-translational modification of Tyr126 that generates the intrinsic DOPA cofactor is NrdI and O_2_ dependent, and may potentially also require other factors, including metals. Still, even if this would be the case, it would require only catalytic amounts of metal in relation to the R2 protein, drastically reducing the amount of metal required for the RNR machinery in the organisms encoding this type of RNR. Detailed characterization of the maturation and radical generation process are exciting future prospects.

## Acknowledgments

We wish to thank Prof. James A Imlay for the kind gift of the *E. coli* (Δ*nrdAB*, Δ*nrdEF*) strain, Prof. Eduard Torrents for providing us with the pBAD18 plasmid and Marta Hammerstad for the pET-22b*BcnrdF* plasmid. Financial support to M.H. was provided by the Swedish Research Council (2017-04018), the European Research Council (HIGH-GEAR 724394), and the Knut and Alice Wallenberg Foundation (Wallenberg Academy Fellows (2012.0233 and 2017.0275)), to B.M.S. by the Swedish Research Council (2016-01920), the Swedish Cancer Foundation (CAN 2016/670), and the Wenner-Gren Foundations, to NC by the Australian Research Council (FT140100834).

We would like to thank Diamond Light Source for beamtime (proposals mx11265 and mx15806) and particularly the staff from beamlines I24 and B21.

## Author contributions

B-M.S. and M.H. conceived and led the study. D.L. performed bioinformatics. V.S. did cloning and operon constructs. V.S. and H.L performed protein production, in vivo and in vitro activity assays, TXRF analysis and crystallography. Y.K. M.S. and N.C. performed spectroscopy. M.L. performed SAXS experiments. J.E. and R.M.M.B. performed mass spectrometry. All authors were involved in experiment design and data analysis. M.H. wrote the manuscript with significant input from all authors.

## Author Information

The authors declare no competing financial interests. Correspondence and requests for materials should be addressed to M.H.

## Accession codes

Structures and crystallographic data have been deposited in the protein data bank with PDB id: 6GP2, DOPA-active form and 6GP3, inactive form.

## Materials and methods

### Bioinformatics

NCBI’s RefSeq database was searched with custom NrdF HMMER (*41*) profiles on 17 February 2017 resulting in 4620 sequences. QSK/VPK variants were manually identified. To reduce the number of highly similar sequences in the phylogeny, the sequences were clustered at 0.75 identity using USEARCH (*42*) and poor quality sequences were manually removed. The remaining 181 sequences were aligned with ProbCons (*43*) and reliable columns in the alignment were identified with the BMGE program using the BLOSUM30 matrix (*44*). A maximum likelihood phylogeny was estimated from the alignment with RAxML version 8.2.4 (*45*), using the PROTGAMMAAUTO model with maximum likelihood estimation starting from a population of 750 rapid bootstrap trees. The number of bootstrap trees was determined determined with the MRE-based bootstopping criterium.

### Cloning

Genomic DNA of *Mesoplasma florum* was extracted from a *Mesoplasma florum* L1 (NCTC 11704) culture obtained from the National Collection of Type Cultures operated by Public Health England. The *M. florum nrdF* gene was amplified from *M. florum* genomic DNA by PCR using *MfnrdF-fow* and *MfnrdF-rev* primers and ligated between NheI-BamHI restriction sites of a modified pET-28a plasmid (Novagen), in which the thrombin cleavage site following the N-terminal His6 tag has been replaced by a TEV site. All codons translating to Tryptophan in the *MfnrdF* gene were mutated from TGA to TGG to correct for the codon usage difference between *M. florum* and *E. coli*. The QuikChange Lightning Multi Site-Directed Mutagenesis Kit from Agilent Technologies, using *MfnrdF-mut1*, *MfnrdF-mut2*, *MfnrdF-mut3* and *MfnrdF-mut4* primers (Table S6), as per the protocol suggested by the manufactures, were used for these changes. Similarly, the pET28*MfnrdI* plasmid was generated by ligating the *MfnrdI* PCR amplicon resulting from *MfnrdI-Fow* and *MfnrdI-Rev* primers, into the modified pET-28a plasmid between NdeI-BamHI restriction sites. A synthetic gene coding for *M. florum nrdE* with a N-terminal His6 tag and a TEV cleavable site, into a pET-21a vector in between the restriction sites NdeI-EcoRI and codon optimized for overexpression in *E. coli* was ordered from GenScript (pET21*MfnrdE*). To generate the pET28*MfnrdFIE* plasmid the *MfnrdF, MfnrdI* and *MfnrdE* genes were individually amplified with primer pairs of *MfnrdF-Fow-MfnrdF-Rev, MfnrdI2-Fow-MfnrdI2-Rev* and *MfnrdE-Fow-MfnrdE-Rev* and double digested by restriction enzymes pairs of NheI-BamHI, BamHI-SacI and Sacl-Sall respectively. The amplicons were then ligated into the NheI and SalI restriction site of a modified pET-28a plasmid with *MfnrdF* placed with a N-terminal TEV cleavable HIS-tag and ribosome binding sites placed ahead of both *MfNrdI* and *MfNrdE* genes (Fig. S11). This entire operon assembly was double digested from pET28*MfnrdFIE* and ligated between KpnI and SalI restriction digestion sites of pBAD18 to generate the pBAD18*MfnrdFIE* plasmid. *pET28MfnrdFI* was prepared similarly by double digesting pET28*MfnrdFIE* with NheI and SacI and ligating it into a modified pET-28a plasmid. All of the above-mentioned plasmids were sequenced to check for any unintended mutations.

### In vivo *studies*

A 4 ml LB (ForMedium) culture of *E. coli* double knockout JEM164 (Δ*nrdAB, ΔnrdEF~srlD::Tn10*) supplemented with tetracycline was grown in an anerobic glovebox (MBraun Unilab Plus SP model) with O_2_ < 10 ppm, washed twice with ice cold water to prepare electro competent cells, transformed with pBAD18*MfnrdFIE* plasmid and plated on a LB-agar media with tetracycline and carbenicillin. Primary cultures of both JEM164 and JEM164-pBAD18*MfnrdFIE* were grown at 37°C overnight anaerobically, supplemented with respective antibiotics. The cultures were removed from the glovebox and inoculated into aerobic LB media supplemented with tetracycline and 0.1% L-arabinose for JEM164 and tetracycline, carbenicillin and either 0.1% D-glucose or 0.1% L-arabinose for JEM164-pBAD18*MfnrdFIE* and grown for 48 hours at 37°C in a bench-top bioreactor system (Harbinger). Tetracycline was used at a concentration of 12.5 μg/ml and carbenicillin was used at a concentration of 50 μg/ml for all of the above-mentioned cultures.

### Protein expression in aerobic conditions

*E. coli* BL21(DE3) (NEB) carrying the plasmid pRARE *camR* (Novagen) were transformed with each of the pET28*MfnrdF*, pET28*MfinrdI*, pET28*MfnrdFI* and pET28*MfnrdFIE* plasmids individually. Glycerol stocks of the transformed cells were flash-frozen and stored at −80°C. All cultures were grown in LB media at 37° C, supplemented with 50 μg/ml kanamycin and 25 μg/ml chloramphenicol, in a bench-top bioreactor system (Harbinger) until an OD600 of ~ 0.7 was reached followed by induction with 0.5 mM isopropyl β-D-1-thiogalactopyranoside (IPTG). The cultures were then allowed to grow overnight at room temperature. pET21*MfnrdE*-BL21(DE3) was grown similarly, but with an antibiotic selection of 50 μg/ml carbenicillin instead. Post-expression the cells were harvested by centrifugation and stored at −20°C until further use.

### Protein purification

The cell pellets were thawed and re-suspended in lysis buffer (25 mM HEPES-Na pH 7, 20 mM imidazole and 300 mM NaCl) and lysed using a high-pressure homogenizer (EmulsiFlex-C3). The un-lysed cells and cell debris was pelleted by centrifugation and the clear supernatant was applied to a lysis buffer-equilibrated gravity-flow Ni-NTA agarose resin (Protino) column and washed with lysis buffer of 40 mM imadizole concentration. The bound protein was eluted with elution buffer (25 mM HEPES-Na pH 7, 250 mM imidazole and 300 mM NaCl), concentrated and loaded onto a HiLoad 16/60 Superdex 200 size exclusion column (GE Healthcare) attached to an ÄktaPrime Plus (GE Healthcare) equilibrated with SEC buffer (25 mM HEPES-Na pH 7 and 50 mM NaCl). The fractions corresponding to *Mf*R2 was pooled and treated to TEV protease cleavage overnight, at a molar ratio of one TEV protease for every 50 MfR2 monomers. TEV protease and any un-cleaved HIS-tagged *Mf*R2 was removed by performing an additional reverse Ni-NTA step. The purity of the protein was evaluated using SDS-PAGE, the protein was then concentrated using a Vivaspin 20 centrifugal concentrators 30,000 Da molecular weight cut-off polyethersulfone membrane (Sartorius) to a desired concentration. All of the above purifications steps were performed at 4° C. The purified *Mf*R2 protein was then aliquoted, flash-frozen in liquid nitrogen and stored at −80°C. *Mf*NrdI and *Mf*R1 were purified similarly but with a buffer system of 25 mM Tris-HCl pH 8 instead of 25 mM HEPES-Na pH 7.

### Anaerobic MfR2 production and purification

The glycerol stock of *E. coli* transformed with the pET28*MfnrdFIE* was used to inoculate a primary culture of LB media supplemented with 50 μg/ml kanamycin and 25 μg/ml chloramphenicol and grown in aerobic conditions at 37°C overnight. All following steps of protein production and purification were carried out anaerobically in an anaerobic chamber. The primary culture was transferred into the anaerobic chamber and used to inoculate at 2% (v/v) a secondary culture of TB media (ForMedium) deoxygenated by N_2_-saturation and supplemented with 50 μg/ml kanamycin and 25 μg/ml chloramphenicol, and grown anaerobically at 37°C until an OD600 of 1.1 was reached. This secondary culture was then used to inoculate deoxygenated TB media supplemented with 50 μg/ml kanamycin and 25 μg/ml chloramphenicol. As the cell culture reached an OD_600_ of 0.7, overexpression was induced with 0.5 mM IPTG and allowed to grow overnight at 37°C, the cells were then harvested by centrifugation. In order to extract the soluble fraction, the cell pellet was resuspended at room temperature using 5 ml of deoxygenated BugBuster Protein Extraction Reagent (Novagen) per gram of wet cell paste. The cell suspension was stirred for 40 min at room temperature. The insoluble cell debris were removed by centrifugation at 13,000 × g for 20 min. From the supernatant, the *Mf*R2 protein was purified, as described in the aerobic procedure, using deoxygenated buffers. Instead of size exclusion chromatography, a HiTrap Desalting column (GE Healthcare) was used to remove the imidazole. The protein was then subjected to TEV protease treatment at a molar ratio of one TEV protease for every 10 *Mf*R2 monomers at room temperature for 1 h. TEV protease and any un-cleaved HIS-tagged *Mf*R2 was removed by passing the sample over a Ni-NTA agarose (Protino) gravity flow column equilibrated in the SEC buffer (25 mM HEPES-Na pH 7.0, 50 mM NaCl). The flow-through containing pure cleaved *Mf*R2 was concentrated to a desired concentration, aliquoted, flash-frozen in liquid nitrogen and stored at −80 °C.

### *In vitro* activity

The *in-vitro M. florum* R1-R2 catalyzed reduction of CDP to deoxy-CDP was measured in a HPLC system (Agilent 1260 Infinity) using a Waters Symmetry C18 column. A 50 μ1 reaction mixture consists of 5 μM *Mf*R1 protein, 5 μM *Mf*R2 protein, 50 mM of Tris-HCl pH 8.0, 20 mM MgCl_2_, 50 mM DTT and 0.5 mM of dATP. The reaction was started with the addition of 2 mM CDP and allowed to run at room temperature for 30-60 minutes, then quenched with 50 μL of Methanol. Additional 200 μL of Milli-Q water was added to the reaction mixture. The concentration of the deoxy-CDP product was measured as described previously(*18, 46*). All of the *in-vitro* RNR enzymatic reactions were performed with four replicates.

### Crystallization

*M. florum* R2 was crystallized by sitting drop vapor diffusion method at a protein concentration of 25 mg/ml. A Mosquito nanoliter pipetting robot (TTP Labtech) was used to set up MRC 2-well crystallization plates (Swissci) with 200:200 nl protein to reservoir volume ratios and reservoir volume of 50 μl. Initial crystallization hits were obtained in condition number C4 of the PEG/Ion HT crystallization screen (Hampton Research). The crystallization condition was further optimized using the Additive screen HT (Hampton Research) with condition number B3 producing the best hits. The crystallization condition is as follows: 100-200 mM Calcium acetate,100 mM Ammonium sulfate and 15-20% PEG 3350, with cuboidal crystals appearing within 2-3 days of incubation at room temperature. Reservoir solution supplemented with 20% glycerol was used as cryoprotectant. Crystals were flash-frozen in liquid nitrogen before data collection

### Data collection and structure determination

X-ray diffraction data were collected at beamline I24 of the Diamond Light Source (Oxfordshire, United Kingdom) at a wavelength of 0.96859 Å at 100 K. Data reduction was carried out using XDS (*47*).

*Mf*R2 crystal structures were solved using PHASER (*48*) by molecular replacement using the atomic coordinates of the R2 protein from *Corynebacterium ammoniagenes* (PDB: 3dhz) (*49*) as a starting model. A well-contrasted solution was obtained with 2 molecules per asymmetric unit in the space group C2 for all the datasets. Crystallographic refinement was performed using PHENIX (*50*) applying anisotropic B-factor and TLS, and was similarly but independently conducted for the 3 models. The 3D models were examined and modified using the program COOT (*51*) and validated using MolProbity (*52*). The core root-mean-square deviation values between structures were calculated by the Secondary-Structure Matching (SSM) tool (*53*). Table S1 was generated with phenix.table_one and lists the crystallographic statistics in which the test set represents 5% of the reflections. The Ramachandran statistics are: favored 98.5%, 99.2% and 99.5% for the active, inactive aerobic and inactive anaerobic *Mf*R2 crystal structures, respectively. Ramachandran outliers were 0.0% in all cases. Figure 2 was prepared using the PyMOL Molecular Graphics System, Version 2.0 Schrödinger, LLC.

For anomalous signal analysis, data were processed with XDS keeping the Friedel pair reflections unmerged (FRIEDEL’S_LAW=FALSE) to preserve any anomalous contributions. Anomalous difference maps were calculated with PHENIX.

### EPR sample preparation

Class Ia *E. coli nrdB* was PCR amplified using primers pairs of *EcnrdB*-Fow and *EcnrdB*-Rev from genomic DNA of *E. coli* BL21 (DE3) (Novagen). The PCR amplicon was then double digested with restriction enzymes pair of NheI-BamHI and ligated into a modified pET-28a plasmid. Class Ib *Bacillus cereus nrdF* gene was restriction digested out of pET-22b*BcnrdF* using NdeI-HindIII restriction enzymes and ligated into a modified pET-28a plasmid. The resulting plasmids were sequenced for any unintended mutations. The *E. coli R2a* and *B. cereus R2b* proteins were produced and purified using protocols as described earlier. A few modifications to the protocols were made in order to obtain metal-free R2 proteins i.e., addition of 0.5 mM of ethylenediaminetetraacetic acid (EDTA) to the growth cultures before induction of overexpression with 0.5 mM IPTG and addition of 0.5 mM of EDTA to the lysis buffer. The metal-free *E. coli* R2 and *B. cereus* R2 proteins were then metal loaded with (NH_4_)_2_Fe(SO_4_)_2_ solution with a final concentration of 1.5 molar equivalent of Fe(II) per monomer and left to incubate at room temperature for 60 minutes under aerobic conditions. The R2 proteins were later diluted with 100% glycerol to a final protein concentration of 0.575 mM and 0.695 mM respectively. Active *M. florum* R2 CW-X band EPR sample was also diluted with 100% glycerol to a final protein concentration of 0.675 mM.

### CW-X band EPR measurements

X-Band CW-EPR measurements were performed at 70 K using a Bruker E500 spectrometer equipped with a Bruker ER 4116DM TE_102_/TE_012_ resonator, Oxford Instruments ESR 935 cryostat and ITC-503 temperature controller. Microwave power was 20 μW and magnetic field modulation amplitude was 0.5 G. The static magnetic field was corrected (for X- and Q-band measurements) using a Bruker ER 035M NMR Gaussmeter. For measurements at 103K the temperature was controlled by flowing gas of known temperature over the sample in a dewar in the cavity. For spectra at 298 K the sample was contained in a capillary. Quantifications were performed by comparing double integrals of the samples with that a standard Cu^2+^/EDTA (1 mM and 10 mM, respectively) sample at non-saturating conditions.

### Microwave saturation measurements

The amplitude of the first derivative of an EPR signal is a function of the applied microwave power (P). Plotting the normalized intensity of the EPR signal X’=(Y’/P^1/2^)/(Y’o/P0^1/2^) as a function of P in the logarithmic form allows determination of the half-saturation power (P_1/2_) (*30*). Under non-saturating conditions X’ equals 1. As the signal starts to saturate, X’ decreases.

The half-saturation power is calculated from the equation X’= 1/(1+P/P_1/2_)^1/2^ for an inhomogeneously broadened signal. The value of P_1/2_ is affected by the magnetic environment of the studied radical. The temperature dependence of P_1/2_ can provide information on an adjacent metal center, or it may reflect the absence of metals in the vicinity of the radical. R2Tyr^•^ with adjacent diiron centers have P_1/2_ values that are considerably higher than those for isolated Tyr^•^ radicals at temperatures above 20-30 K (*30*).

### Pulse-Q band EPR measurements

Q-band pulse-EPR measurements were performed in the temperature range of 50 to 70 K using a Bruker ELEXSYS E580 Q-band pulse-EPR spectrometer, equipped with a homebuilt TE_011_ microwave cavity(*54*) Oxford-CF935 liquid helium cryostat and an ITC-502S temperature controller. Electron spin echo-detected (ESE) field-swept spectra were measured using the pulse sequence: t_p_-*τ*-2t_p_-*τ*-echo. The length of the π/2 microwave pulse was generally set to t_p_ = 22 ns and the interpulse distance was *τ* = 260 ns. Pseudomodulated EPR spectra were generated by convoluting the original absorption spectra with a Bessel function of the 1^st^ kind. The peak-to-peak amplitude used was 0.8 G. ^1^H-ENDOR spectra were collected using the Davis pulse sequence: t*_inv_* - t*_RF_* - T - t_p_ -*τ*- 2t_p_ -*τ*-echo with an inversion microwave pulse length of t*_inv_* = 140 ns, a radio frequency pulse length of t_RF_ = 20 μs (optimized for ^1^H). The length of the π/2 microwave pulse in the detection sequence was set to t_p_ = 70 ns and the interpulse delays to *T* = 22 μs and *τ* = 400 ns. The RF frequency was swept 40 MHz around the ^1^H-Larmor frequency of about 52 MHz (at 1.2 T) in 100 or 50 kHz steps. The RF amplifier used for ENDOR measurements was ENI 3200L.

### Total-reflection X-ray fluorescence (TXRF) metal quantification

The metal contents of protein and buffer solutions were quantified using TXRF analysis on a Bruker PicoFox S2 instrument. For each solution, measurements on 3 independently prepared samples were carried out. A gallium internal standard at 2 mg/l was added to the samples (v/v 1:1) prior to the measurements. TXRF spectra were analyzed using the software provided with the spectrometer. As a control, the metal-free *E. coli* class Ia R2 protein was Fe-reconstituted by incubation with 2 molar equivalent of Fe(II) per monomer at room temperature for 60 min under aerobic conditions. The protein solution was then applied on a HiTrap Desalting column (GE Healthcare) in order to remove unbound iron, and was concentrated using a Vivaspin centrifugal concentrator, to a concentration similar to the *Mf*R2 sample used for TXRF measurements.

### Small Angle X-Ray Scattering

SAXS measurements were carried out on beamline B21 at the Diamond Light Source at 12.4 keV in the momentum transfer range 0.0038 <q<0.41 Å^−1^ (q = 4π sin (θ)/λ, 2θ is the scattering angle) using a Pilatus 2M hybrid photon-counting detector (Dectris Ltd., Baden, Switzerland). Prior to measurements, samples of *Mf*R2 alone or *Mf*R2 incubated with MfNrdI, with a molar ratio R2:NrdI of 1:1.5, were run on a HiLoad 16/60 Superdex 200 prep grade size exclusion chromatography column (GE Healthcare) equilibrated in the SEC buffer 25 mM HEPES pH 7.0, 50 mM NaCl. Fractions corresponding to*Mf*R2 or to the complex of*Mf*R2-MfNrdI were pooled, diluted to 3 different concentrations with the SEC buffer and flash-frozen until measurements. A volume of 30 μl of either*Mf*R2 alone or*Mf*R2 incubated withMfNrdI was loaded onto the sample capillary using the EMBL Arinax sample handling robot. Each dataset comprised 18 exposures each of 180 seconds. Identical buffer samples were measured before and after each protein measurement and used for background subtraction. Data averaging and subtracting used the data processing tools of the EMBL-Hamburg ATSAS package (*55*). The radius of gyration (Rg) and maximum particle size (Dmax) were determined from tools in the PRIMUS program suite (*56*). Twenty independent *ab initio* models of either *Mf*R2 or the *Mf*R2-NrdI complex were derived from the experimental scattering curves using DAMMIF (*57*). For each protein, the models were aligned, averaged and filtered using the DAMAVER program suite (*58*). Theoretical scattering profiles of the crystal structure of *Mf*R2 and the *Mf*R2-NrdI homology model were calculated and fitted against the experimental data using CRYSOL (*59*). The homology model of the *Mf*R2-NrdI complex was created by superposing the *Mf*R2 dimer crystal structure and an MfNrdI homology model prepared based on the crystal structure of the R2-NrdI complex from *E. coli* (PDB: 3N3A) (*21*).

### Proteolytic digestion and peptide analysis by LC-MS

The protein of interest was cleaned from contaminants in the SEC buffer following a modified version of the SP3 protocol (*60, 61*). Samples were digested and subjected to LC-MS/MS analysis. Unless noted otherwise, all reagents were purchased from Sigma Aldrich.

The SP3 beads stock suspension was prepared freshly prior to sample processing. The two SP3 bead bottles (Sera-Mag Speed beads - carboxylate modified particles, P/N 65152105050250 + P/N 45152105050250 from Thermo Scientific) were shaken gently until the suspension looked homogenous, and then 50 μl from each bottle was taken to one tube. The 100μl of beads was then washed three times with water (in each wash, tubes were taken off the magnetic rack, beads were mixed with 500 μl H2O by pipetting, then placed back in the rack, and after a 30 s delay, the supernatant was removed). Beads were finally resuspended in 500 μl H_2_O after washing. To each aliquot of 30 μl of purified protein solution in SEC buffer at 70 μM protein concentration, 10 μl of SP3 stock suspension was added. MeCN (acetonitrile) was added to reach a final concentration of 50 % (v/v). The reaction was allowed to stand for 20 min. The beads were collected on the magnetic rack and the supernatant was discarded. The beads were washed two times with 70 % (v/v) EtOH and once with MeCN. The beads were then resuspended in 100 μl of digestion buffer containing either i) 1 μg of Chymotrypsin in 50 mM HEPES, pH 8, 10 mM CaCl_2_, or ii) 1 μg of Pepsin in 50 mM HCl, pH 1.37. Samples were digested at 37°C overnight. The beads were removed and where necessary HCl was neutralized with one equivalent of triethylammonium bicarbonate buffer (1M, pH 8.5). From each sample, 8μl was injected to LC-MS/MS analysis on an LTQ-Orbitrap Velos Pro coupled to an Agilent 1200 nano-LC system. Samples were trapped on a Zorbax 300SB-C_18_ column (0.3×5 mm, 5 μm particle size) and separated on a NTCC-360/100-5-153 (100 μm internal diameter, 150 mm long, 5 μm particle size, Nikkyo Technos., Ltd, Tokyo, Japan, (http://www.nikkyo-tec.co.jp) picofrit column using mobile phases A (3% MeCN, 0.1% FA) and B (95% MeCN, 0.1% FA) with a gradient ranging from 3% to 40% B in 45 min at a flowrate of 0.4 μl/min. The LTQ-Orbitrap Velos was operated in a data-dependent manner, selecting up to 3 precursors for sequential fragmentation by both CID and HCD, analyzed in the Orbitrap. The survey scan was performed in the Orbitrap at 30000 resolution (profile mode) from 300-1700 m/z with a max injection time of 200 ms and AGC set to 1 × 10^6^ ions. MS2 scans were acquired at 15 000 resolution in the Orbitrap. Peptides for dissociation were accumulated for a max ion injection time of 500 ms and AGC of 5 × 10^4^ with 35% collision energy. Precursors were isolated with a width of 2 m/z and put on the exclusion list for 30 s after two repetitions. Unassigned charge states were rejected from precursor selection. The raw files were searched against a database only containing the sequence of NrdF R2 protein with Proteome Discoverer 1.4 and the Sequest algorithm (*62*) allowing for a precursor mass tolerance of 15 ppm and fragment mass tolerance of 0.02 Da. Non-enzyme searches were conducted and methionine and tyrosine oxidations were included as variable modifications. For the MODa (*63*) searches, raw files were first converted to mzXML files using msconvert from ProteoWizard (*64*) using vendor peak picking at the MS^2^ level and otherwise standard settings. The mzXML files were searched against the sequence of NrdF with MODa, allowing for an arbitrary number of mods with sizes within the +/− 200 Da interval, and using a fragment mass tolerance of 0.05 Da. Otherwise the same settings as for the Sequest searches were chosen. The high-resolution mode was used.

### Intact protein analysis by LC-MS

A volume of 2 μl of inactive *Mf*R2 and active R2, at a concentration of 70 μM, were diluted in 200μl of LC-MS mobile phase A (3 % MeCN, 0.1% FA). From this, 3 μl was injected in each LC-MS run, which was set up as described in the previous section, but with the following differences: the trapping column was a Zorbax 300SB-C_8_ column (0.3×5mm, 5μm particle size); the gradient went from 3% B to 99% B in 10 min at a flow rate of 0.6 μl/min; the MS only acquired full scans of the 600-4000 m/z high mass range at 100 000 resolution.

The raw MS files were processed with Protein Deconvolution 4.0 (Thermo Scientific), using the ReSpect algorithm therein. Default parameters were used except: the relative abundance threshold was set to 40%, the m/z range set to 800-2400 m/z and the charge state range set to 6-100.

## Supplementary information

**Figure S1.**
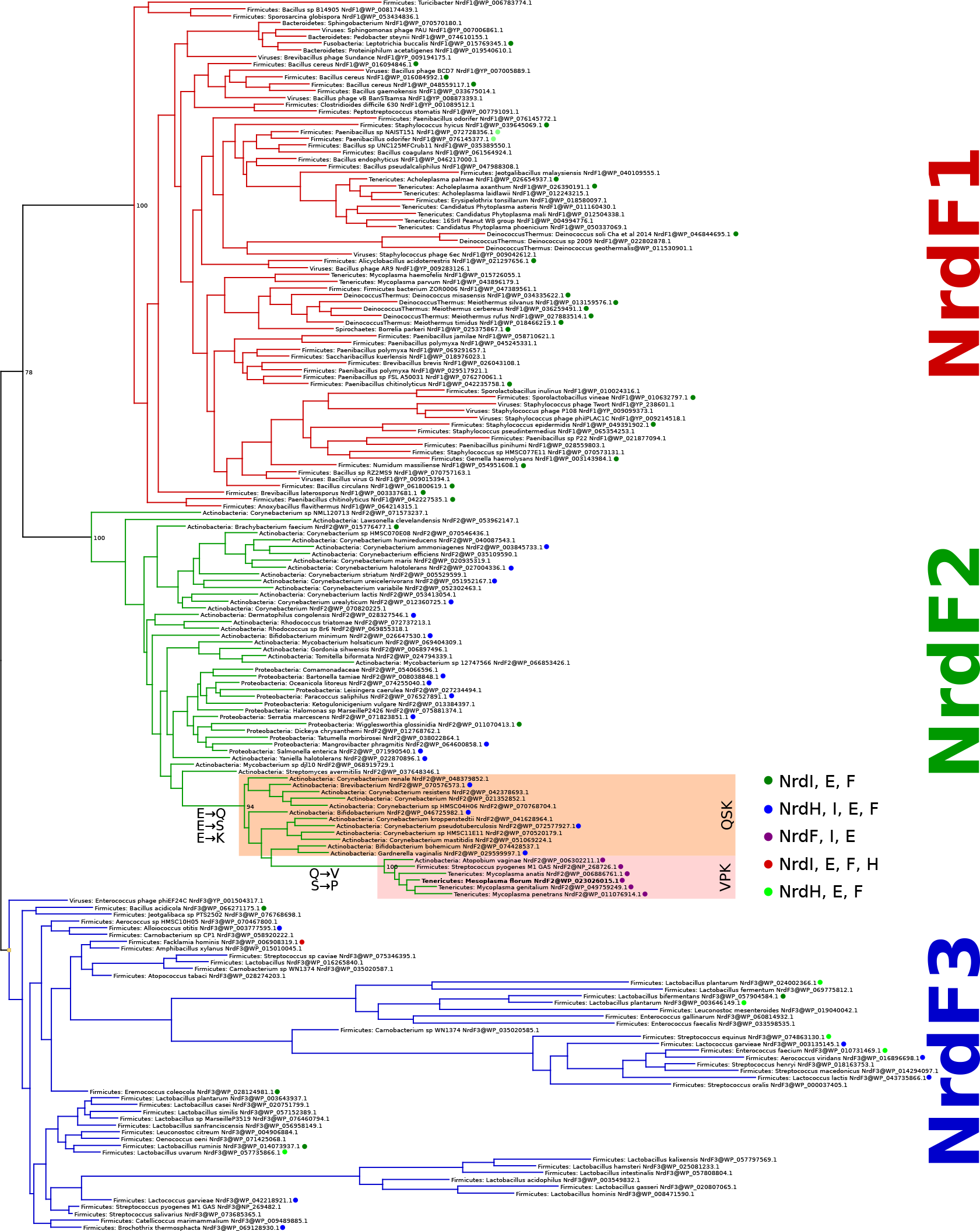
Unrooted maximum likelihood phylogeny of representative NrdF (RNR subclass Ib radical-generating subunit) sequences. All RefSeq NrdF sequences were clustered at 75% identity to reduce redundancy and a maximum likelihood phylogeny was estimated. Sequences with non-canonical amino acids in the positions involved in coordinating the metal center of the enzyme formed a well-supported clan in the NrdF2 group of sequences. We identified two variants, one in which three of the glutamates were replaced by Gln, Ser, and Lys (NrdF2.QSK) and the other in which they were replaced by Val, Pro and Lys (NrdF2.VPK). Both variants thus have a substitution of a Lys for the normally metal-bridging Glu (residue 213 in *M. florum* NrdF2.VPK). Together, the two variants form a well-supported (96% bootstrap support) clan in the phylogeny inside the NrdF2 diversity. The NrdF2.VPK clan appears to be derived from the NrdF2.QSK clan. Behind the sequences in the tree are a set of sequences more than 75% identical to each represented sequence. The VPK and QSK sequences in the phylogeny represent 138 and 182 sequences in RefSeq respectively.

**Figure S2.**
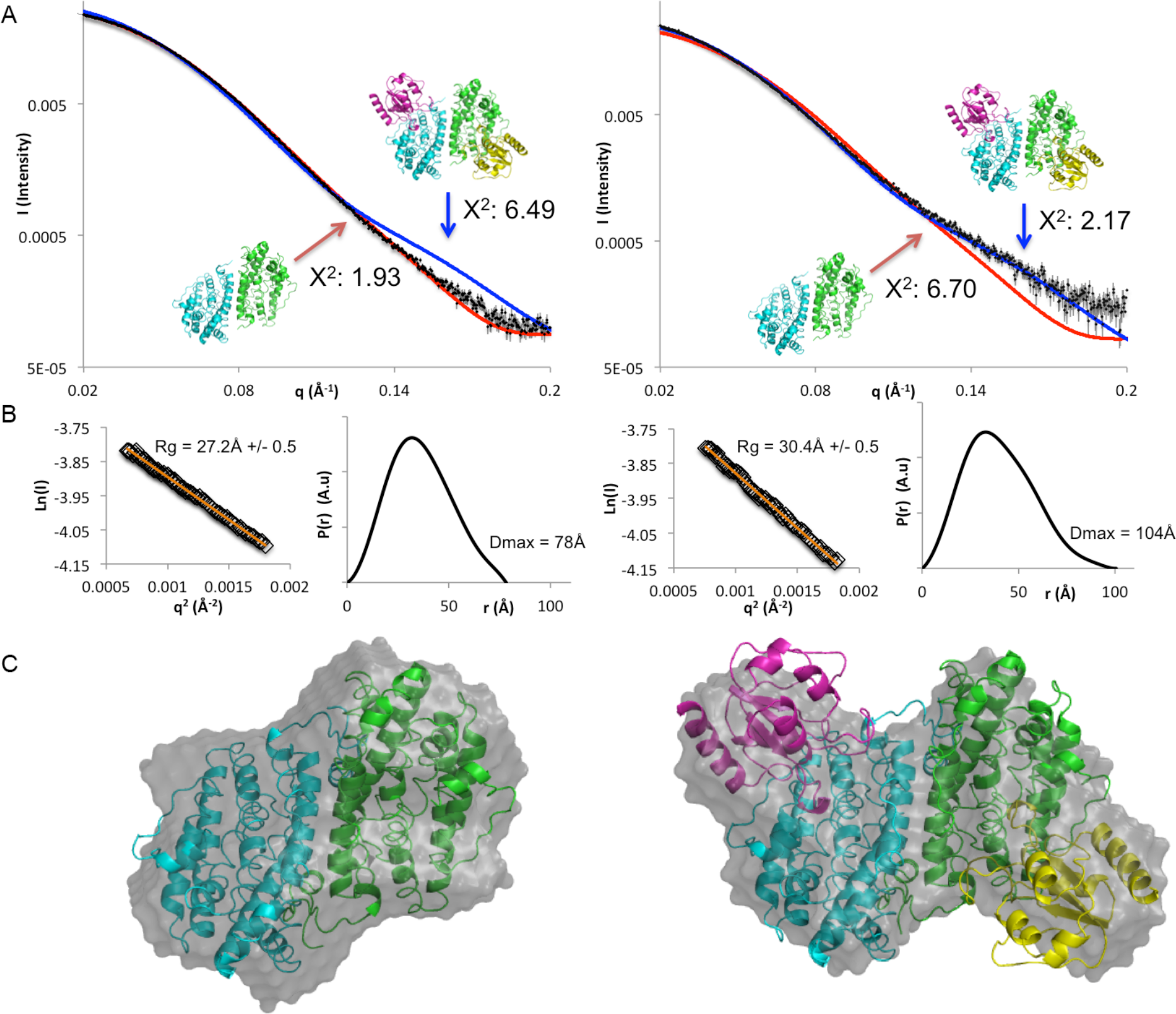
Small angle X-ray scattering characterization of the *Mf*R2-NrdI complex. Solution scattering data for *Mf*R2 (left) and *Mf*R2 incubated with MfNrdI (right). **A)** Experimental solution scattering profiles (black spheres) for *Mf*R2 alone and incubated with MfNrdI superposed with the theoretical scattering profile of the *Mf*R2 crystal structure (red line) and the theoretical scattering profile from the homology model based on the *E. coli* R2-NrdI complex structure (blue line). Theoretical scattering curves and goodness of fit values were calculated by CRYSOL. **B)** Guinier fit and p(r) function of *Mf*R2 alone and incubated with MfNrdI. The fit to the data is shown as an orange line. The shift in invariant parameters R_g_ and D_max_ indicate that an increase in dimensions occurred as *Mf*R2 was incubated with *Mf*NrdI. **C)** *ab initio* models as calculated by DAMMIN of *Mf*R2 alone and with NrdI (grey surface) overlaid with the crystal structure of *Mf*R2 (left) and the homology model based on the *E. coli* R2-NrdI complex structure model (right).

**Table S1.** Data collection and refinement statistics.

|  | Active MfR2 | Inactive MfR2 (aerobic) | Inactive MfR2 (anaerobic) |
| --- | --- | --- | --- |
| **Data collection** |  |  |  |
| Space group | C2 | C2 | C2 |
| Cell dimensions |  |  |  |
| *a*, *b*, *c* (Å) | 176.16, 53.74, 79.28 | 176.19, 53.43, 79.13 | 176.30 53.51 79.10 |
| $\alpha$, $\beta$, $\gamma$ (°) | 90, 108.58, 90 | 90, 108.53, 90 | 90 108.49 90 |
| Resolution (Å) | 42.73-1.48 (1.53-1.48)* | 42.71-1.23 (1.27-1.23) | 44.19-1.24 (1.28-1.24) |
| *R*<sub>merge</sub> | 0.066 (1.403) | 0.054 (1.092) | 0.052 (0.951) |
| *I* / σ*I* | 12.67 (1.17) | 15.13 (1.35) | 15.15 (1.50) |
| Completeness (%) | 99.8 (99.7) | 98.9 (97.3) | 97.4 (94.4) |
| Redundancy | 6.6 (6.4) | 6.4 (6.0) | 6.5 (6.2) |
| **Refinement** |  |  |  |
| Resolution (Å) | 42.73-1.43 (1.53-1.43) | 42.71-1.23 (1.27-1.23) | 44.19-1.24 (1.28-1.24) |
| No. reflections | 117311 (11659) | 200307 (19622) | 192936 (18564) |
| *R*<sub>work</sub> / *R*<sub>free</sub> | 0.1543 (0.3738) / 0.1845 (0.4083) | 0.1466 (0.3296) / 0.1688 (0.3434) | 0.1472 (0.3008) / 0.1630 (0.3105) |
| No. atoms | 6137 | 6848 | 6743 |
| Protein | 5610 | 6128 | 6097 |
| Ligand/ion | 2 | 2 | 2 |
| Water | 525 | 718 | 644 |
| *B*-factors | 39.00 | 27.28 | 28.61 |
| Protein | 38.07 | 25.71 | 27.45 |
| Ligand/ion | 29.44 | 20.44 | 22.58 |
| Water | 48.89 | 40.65 | 39.67 |
| R.m.s. deviations |  |  |  |
| Bond lengths (Å) | 0.007 | 0.006 | 0.006 |
| Bond angles (°) | 1.13 | 1.12 | 1.13 |

\*Values in parentheses are for highest-resolution shell.
A single crystal was used for each dataset.

**Figure S3.**
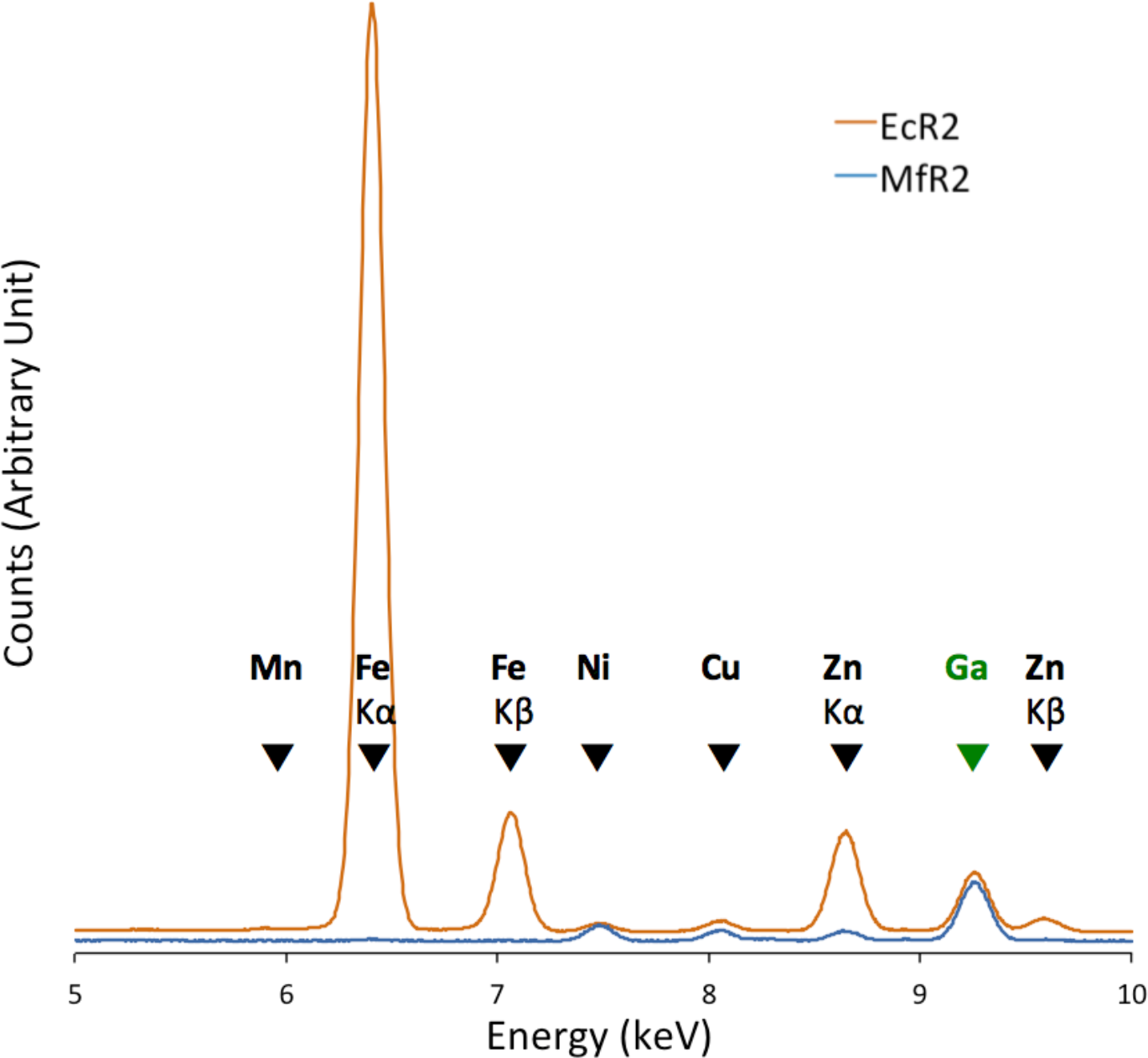
TXRF characterization. Representative TXRF spectra measured for *Mf*R2 (blue, at 664 μM) and Fe-reconstituted class Ia *Ec*R2 (orange, at 635 μM), on the 5 to 10 keV energy range. The spectra have been scaled using the peak size of the Ga internal standard and offset slightly in the Y-direction for clarity. K-level X-ray emission lines are indicated with arrows. For elements, where both Kα and Kβ lines are present, they are specified. Otherwise, arrows indicate Kα lines.

**Table S2.** Total-reflection X-ray fluorescence (TXRF) metal quantification. The concentrations of Mn, Fe, Co, Ni, Cu and Zn were measured in the active *Mf*R2, MfNrdI and *Mf*R1 protein solutions and in their respective buffers, as well as in a solution of *E. coli* class Ia R2 protein reconstituted with Fe *in vitro*. The concentrations and standard deviations reported are averages of measurements on 3 independently prepared samples for each sample. The concentrations were converted to metal to protein molar ratio. The measurements show that none of the *Mf*RNR proteins contain a significant amount of metal as opposed to *Ec*R2a which as expected contains on the order of 2 metal ions per monomer also after a desalting step. Buffer 1 is the buffer system used for *Mf*R2, i.e. 25 mM HEPES-Na pH 7, 50 mM NaCl. Buffer 2 is the buffer system used for *Mf*R1 and *Mf*NrdI, i.e. 25 mM Tris-HCl pH 8, 50 mM NaCl.

|  |  | Mn | Fe | Co | Ni | Cu | Zn | Cumulative |
| --- | --- | --- | --- | --- | --- | --- | --- | --- |
| ***MfR2***<br>**active** | Conc. (μM) | 0,24 ± 0.21 | 2.29 ± 1.02 | n.d.* | 11,28 ± 0.27 | 6,57 ± 0.26 | 4.71 ± 0.33 | - |
|  | mol/mol<br>metal/protein | 0.04% | 0.35% | - | 1.70% | 0.99% | 0.71% | 3.78% |
| ***MfNrdI*** | Conc. (μM) | 0.28 ± 0.24 | 6.84 ± 1.76 | n.d.* | 3.57 ± 0.38 | 6.47 ± 0.96 | 12.20 ± 1.82 | - |
|  | mol/mol<br>metal/protein | 0.03% | 0.76% | - | 0.39% | 0.71% | 1.35% | 3.24% |
| ***MfR1*** | Conc. (μM) | 1.50 ± 0.21 | 2.51 ± 0.19 | n.d.* | 10.50 ± 0.28 | 4.11 ± 0.04 | 13.08 ± 0.12 | - |
|  | mol/mol<br>metal/protein | 0.39% | 0.65% | - | 2.73% | 1.07% | 3.41% | 8.26% |
| ***EcR2a*** | Conc. (μM) | 1.96 ± 0.11 | 1055.79 ± 40.08 | n.d.* | 5.44 ± 0.22 | 6.57 ± 0.09 | 56.97 ± 2.13 | - |
|  | mol/mol<br>metal/protein | 0.31% | 166.27% | - | 0.86% | 1.03% | 8.97% | 177.44% |
| **Buffer**<br>**1** | Conc. (μM) | 0.15 ± 0.15 | 0.12 ± 0.11 | n.d.* | n.d.* | 0.04 ± 0.03 | 0.23 ± 0.02 | - |
| **Buffer**<br>**2** | Conc. (μM) | 0.21 ± 0.25 | 0.19 ± 0.17 | n.d.* | 0.04 ± 0.07 | 0.13 ± 0.05 | 0.20 ± 0.05 | - |

\* n.d. = not detected

**Figure S4.**
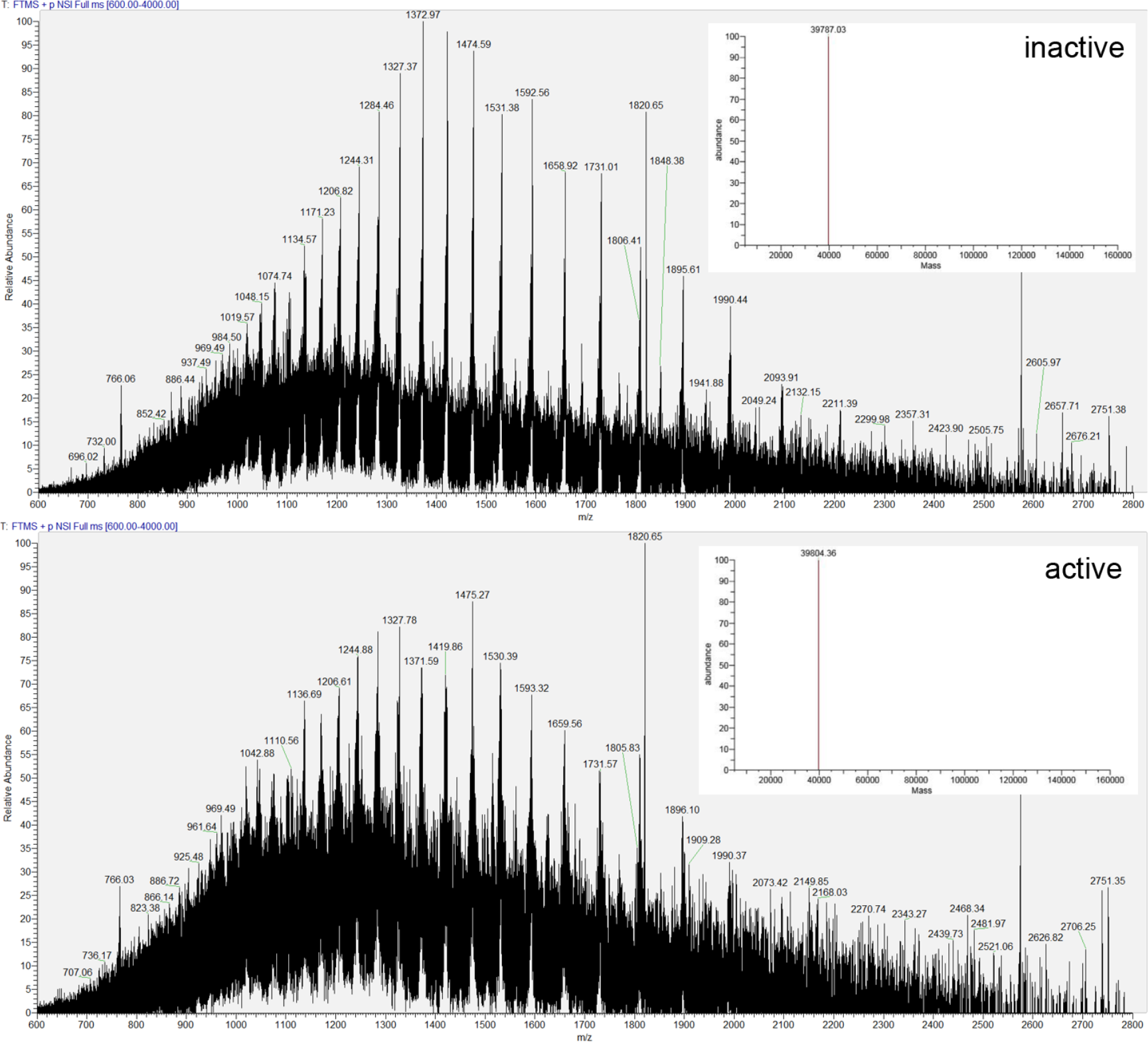
Mass spectrometrometric characterization of intact proteins. Intact protein mass spectra obtained from purified NrdF R2 proteins. Inactive protein (above) and active protein (below). Insets represent the decharged and deisotoped mass as calculated by the program Protein Deconvolution. Mass estimation errors (standard deviations) are 1.4 Da for “inactive” and 2 Da for “active”. The results show that the active protein is 17 ± 2 Da heavier than the inactive *Mf*R2

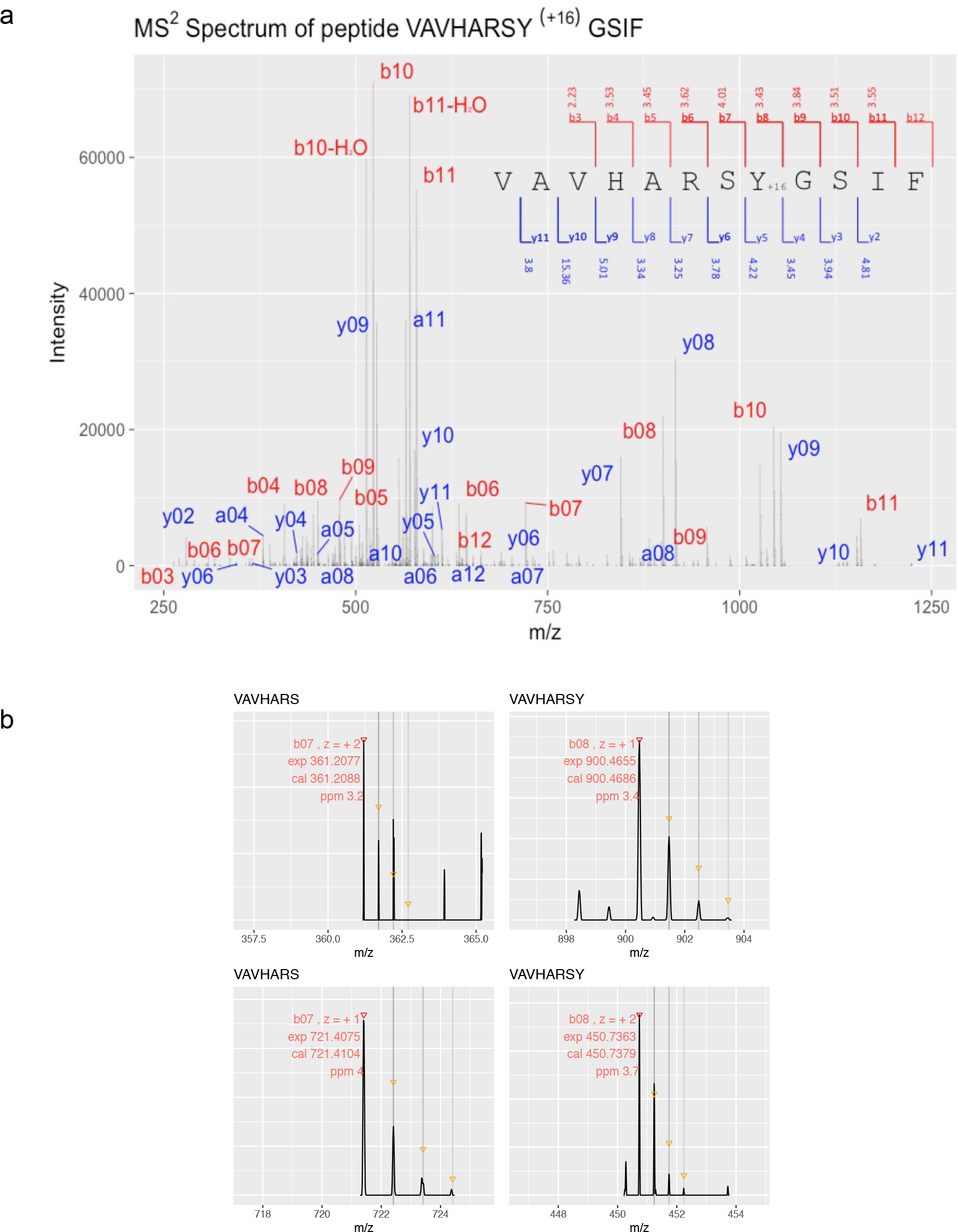

**Figure S5.**
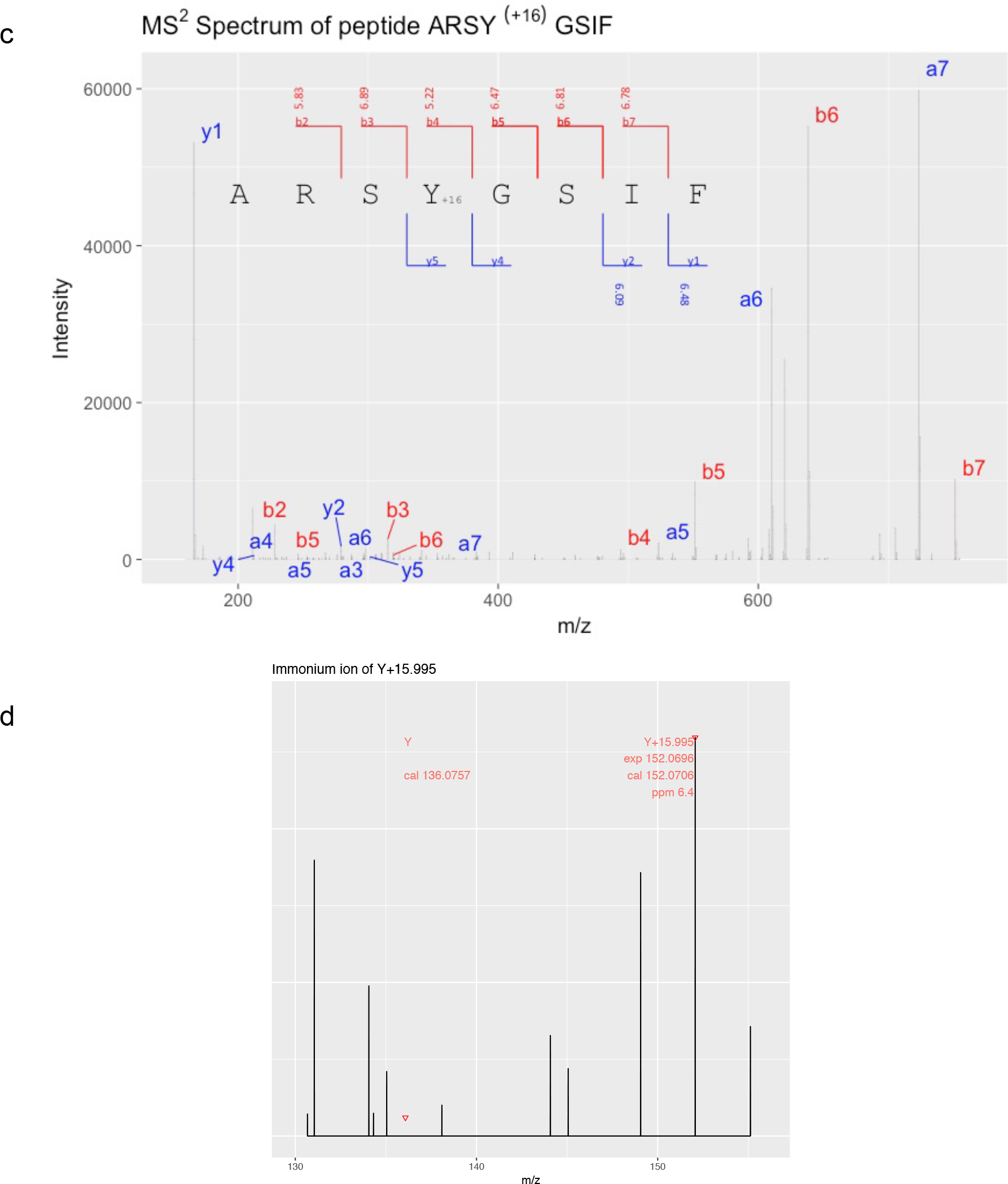
MS2 fragmentation spectra of peptides with oxidized tyrosine. **a)** Annotated MS2 fragmentation spectra and respective theoretical fragment ion tables of the doubly charged precursor ion 661.8458 m/z corresponding to peptide VAVHARSY(+15.995)GSIF, and c) the doubly charged precursor ion 458.7279 m/z corresponding to peptide ARSY(+15.995)GSIF, both with the oxidized (+16) Y126 residue. The peptides shown in **a)** and **c)** were obtained by proteolytic digestion of the active form of the NrdF protein with chymotrypsin and pepsin, respectively. The mass error is typically less than 0.01 m/z, in accordance with the high resolution used (15 000). Errors in ppm are indicated for the corresponding fragment ions when detected. Among the fragment ions observed, the most relevant are the b7 and b8 ions for peptide **a)**. The experimental m/z values, the annotation, theoretical m/z values and ppm errors are shown in **b)**, including the peaks for the corresponding isotope envelope. In d) the Y(+O) immonium ion for **c)** is shown, which demonstrate that Tyr126 is modified by a mass of +15.995 and the absence of the corresponding immonium ion for the unmodified Y.

**Figure S6.**
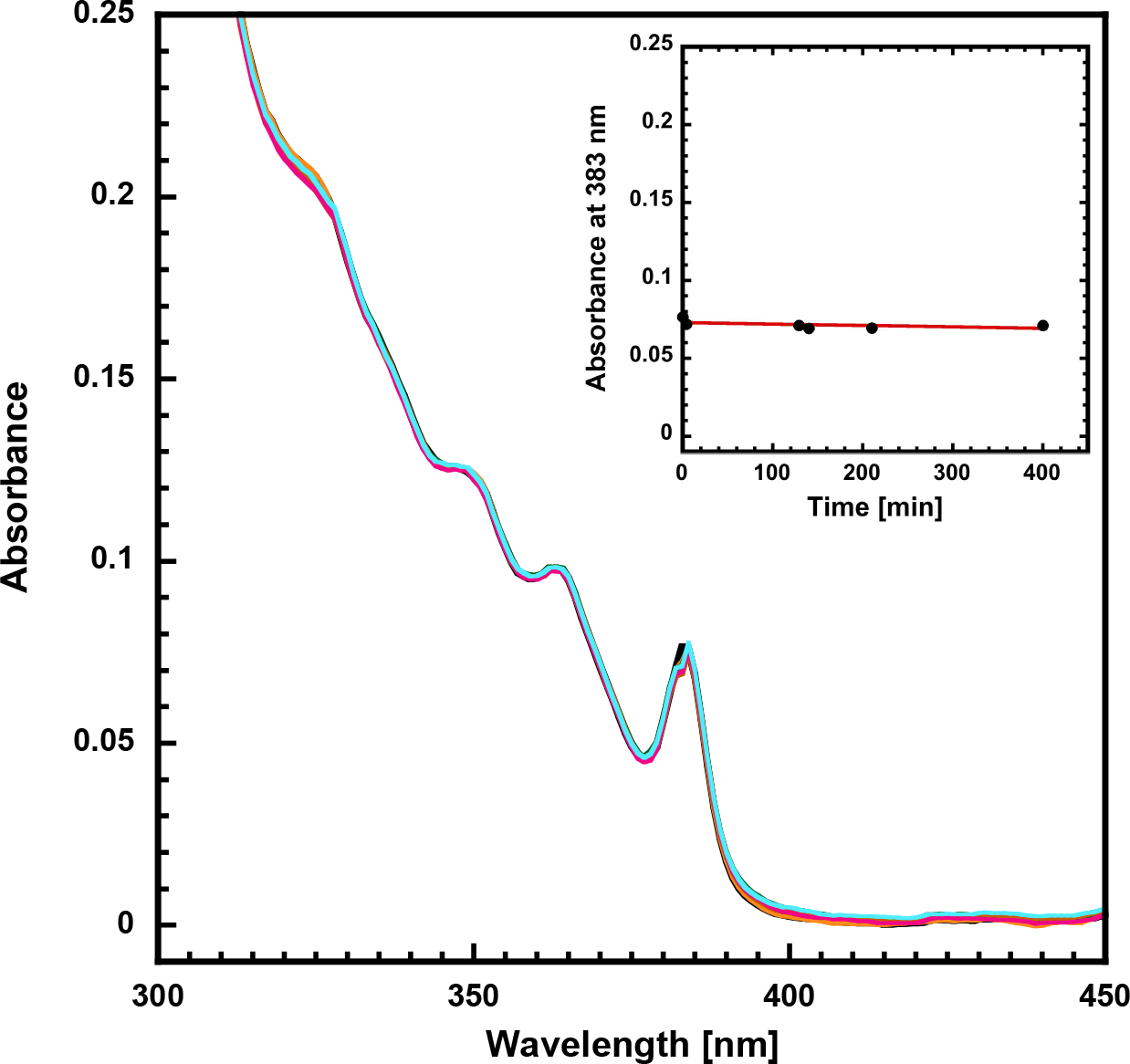
Radical stability. Superimposed UV/vis spectra at timepoints between 0 and 400 minutes. Inset, absorbance at 383 nm at 0, 4, 129, 140, 210 and 400 minutes.

## Supporting information text, EPR characterization

### General description of aromatic free radicals

The lineshape of an EPR signal of a radical the interplay of two magnetic interactions: i) the interaction of the unpaired electron spin with the applied magnetic field (g-value or chemical shift); ii) and the interaction of the unpaired electron spin with local magnetic nuclei via the hyperfine interaction. The g-value defines where the EPR line is observed while the hyperfine interaction leads to a splitting of the EPR line. Hyperfine splittings are additive, with all contributing to the observed spectral width. When measured in the solid state (frozen solution) the g value for each orientation of the molecule relative to the magnetic field axis can be different, with a corresponding different hyperfine splitting. This leads to a broadening of the EPR spectrum and a loss of resolution owing to the overlay of multiple splitting patterns.

The EPR spectra of aromatic free radicals, such as a phenyl radical are complicated owing to the delocalization of the unpaired electron spin across the whole molecule. The unpaired spin resides in the p_z_ orbitals of the carbon atoms which make up the ring, aligned perpendicular to the ring plane. While these orbitals have no direct overlap with the protons of the ring, the exchange interaction of the unpaired spin in the p_z_ orbitals with the paired electron spins of the C-H sigma bonds leads to polarization these bond and thus a hyperfine coupling between ^1^H ring nuclei and the unpaired electron spin(*65*) In addition to the ring protons, the unpaired electron of the conjugated ring will also couple to protons of substituent groups. These coupling are typically larger than those of the ring, as the unpaired electron directly interacts with the protons of the substituent via hyperpolarization (direct overlap)(*65*).

In a phenyl radical the pz orbital of each carbon shares the unpaired spin equally (17%). For a substituted ring this is not the case. For example, for a phenoxy radical it is estimated that the oxy group and the carbon which it is attached to carries slightly higher unpaired spin density of 20-25%, with consequently lower spin densities at the ortho (C2, C6) and particularly the meta (C3, C5) carbon positions of the conjugated ring. Interestingly, it is the para position (C4 on the opposite side of the ring to the phenoxyl group) that carries the highest spin density of 40%.

The magnitude of the ^1^H couplings correlates with the unpaired spin density of the p_z_ orbital of the carbon it is attached to. The approximate magnitude of the coupling is give McConnell’s relation (*65*)

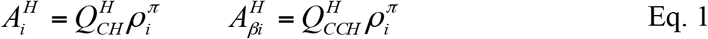

Where Q_CH_ and Q_CCH_ are constant with assigned values of −2.37 mT and 2.72 mT, respectively.

In solution, free rotation about the C-C bond axis of alkyl substituent groups, results in the proton couplings being equivalent. This equivalence is lost in the solid state e.g. a tyrosyl radical embedded in a protein, in which the alkyl protons have a fixed position relative to the ring plane.

**Fig. S7.**
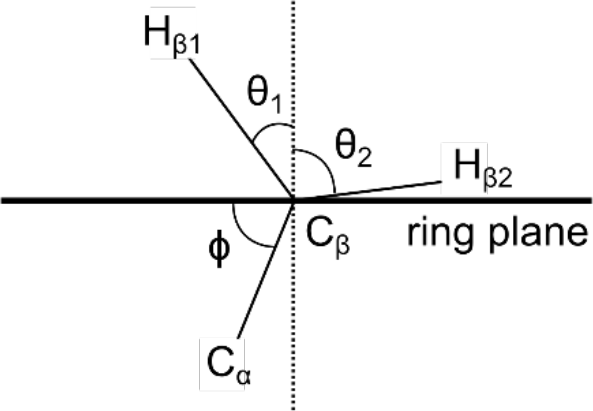
Inferred orientation (θ_1_, θ_2_) of the C_β_ protons relative to the phenoxyl radical ring plane as determined by the dihedral angle (ϕ) between the ring plane (C4) and C_α_.

In this instance the hyperfine coupling is

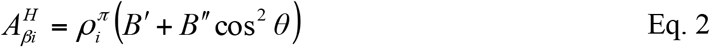

Where *B′* and *B′* are constants and theta is the angle between the proton and a plane normal to the ring plane. *B′* is small and usually ignored. *B′* is 5.8 mT.

### The spin Hamiltonian formalism

A basis set that describes the radical spin manifold can be built from the product of the eigenstates of the interacting electron (*S* = 1/2) and nuclear (^1^H, *I* = 1/2) spins:

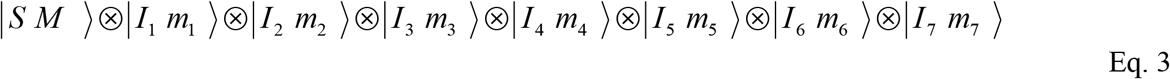

Here, *S* refers to the electronic spin state, M refers to the electronic magnetic sublevel, *I_i_* refers to the nuclear spin state of ^1^H, and m*_i_*, refers to the nuclear magnetic sublevels of each ^1^H. *S* and *I_i_* take the value of ½ and M and m_i_ take the values ±1/2. The spin Hamiltonian that describes the spin manifold is:

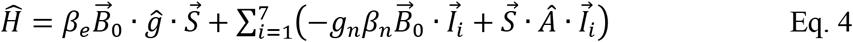

It contains (i) an electronic Zeeman term describing the unpaired electrons interaction with the applied magnetic field (g*_i_*), (iv) a nuclear Zeeman term for each ^1^H nucleus and the applied magnetic field, and (iii) an electron-nuclear hyperfine term for each ^1^H nucleus describing the magnetic interaction between the unpaired electron and each nucleus. Spectral simulations were performed numerically using the EasySpin package (**66, 67**) in MATLAB.

### Description of experimental data and simulated spin Hamiltonian parameters

The EPR spectrum is centered at approximately g = 2.0044. Its spectral width at X- and Q-band is 2.4 and 5.0 mT respectively. The increase in width is due to the anisotropy of the g-tensor. This also gives rise to the asymmetric hyperfine pattern at both X and Q-band. At X-band at least two hyperfine splittings are observed of the order of 1 mT (28 MHz) and 3.6 mT (10 MHz). By way of comparison a typical tyrosine radical always displays at least three hyperfine splitting/coupling in excess of 15 MHz, explaining why the *Mf*R2 radical signal is significantly narrower.

Seven hyperfine couplings were used to simulate the set of EPR and ENDOR spectra collected across the signal. These data constrain the magnitude and tensor symmetry of the three largest hyperfine couplings, although it is noted that the smallest component of tensors A_2_ and A_3_ (i.e. y-component) is unresolved due to spectral congestion. The largest hyperfine coupling is virtually isotropic (axial) and of the order of 30 MHz. It is assigned to one of the protons on C_β_. It gives rise to the broadest ENDOR lines. It is not clear why this is the case. It may reflect local heterogeneity. No second C_β_ proton is observed.

The next three hyperfine couplings are rhombic. For the first two (A2 and A3) A_y_ the unique tensor component. Their tensor structure is very similar to that of protons located at the meta position of a tyrosyl radical. As such they are assigned to two, non-equivalent ring protons. Importantly, the coupling those are systematically smaller (35%) than that seen for a tyrosyl radicals. The third coupling (A4) is less well defined. Its unique tensor component can be assigned to either A_x_ or A_z_. It bears some similarity to that of proton located at the ortho position of a tyrosyl radical and is approximately the same magnitude i.e. 4.5 MHz. The remaining fitted couplings are all dipolar, representing more distant protons in the vicinity of the radical.

**Fig. S8.**
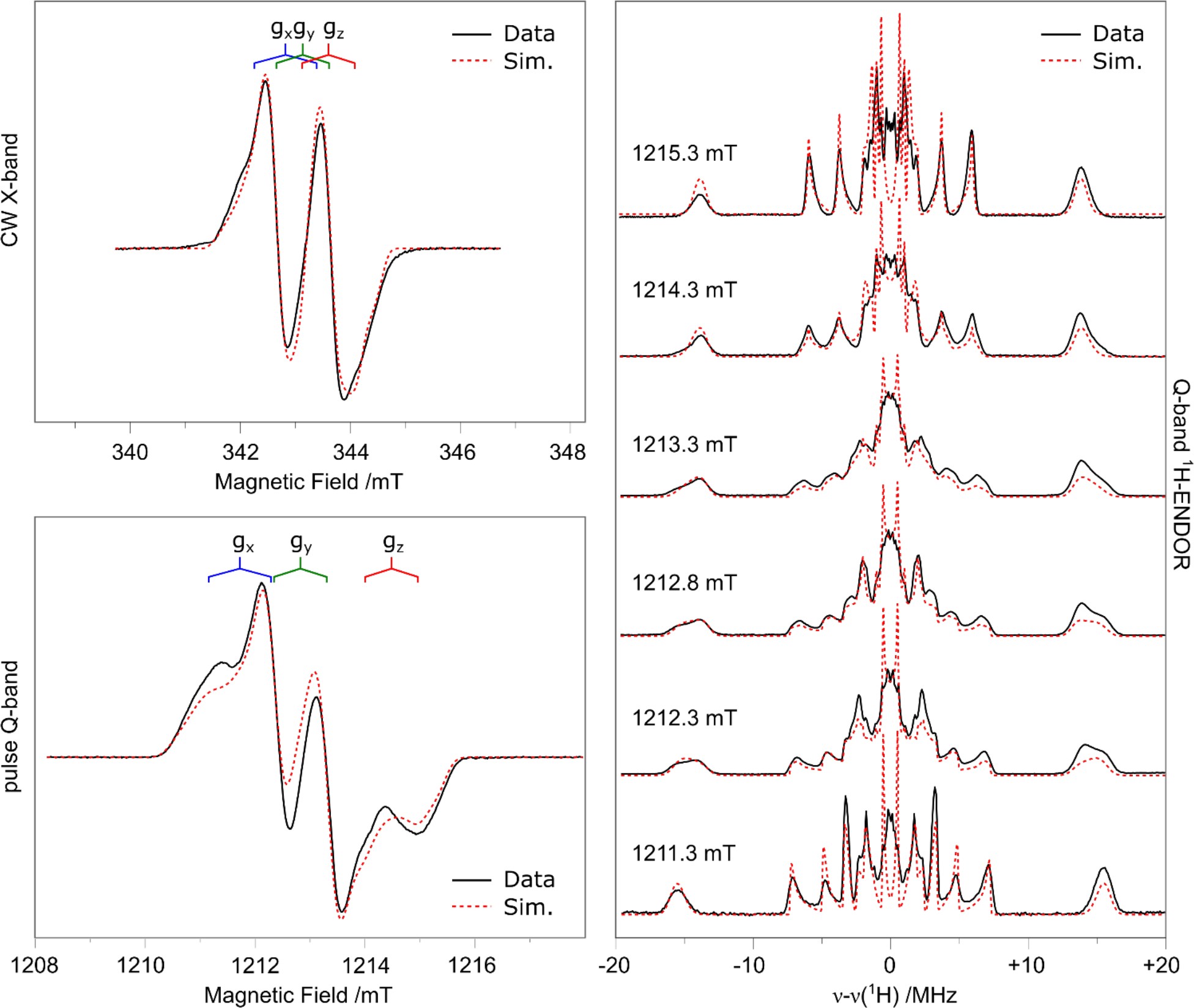
Full mulifrequency (X-/Q-band) EPR dataset and corresponding field dependent Q-band ENDOR spectra. Experimental parameters are listed in the materials and methods section. The red dashed lines represent a simultaneous simulation of all datasets using the spin Hamiltonian formalism. Simulation parameters are listed in Table S2.

**Table S3.** Fitted spin Hamiltonian Parameters.

|  | g-tensor |  |  |  |  |  |
| --- | --- | --- | --- | --- | --- | --- |
|  | x | y | z | iso | aniso |  |
| g | 2.0065 | 2.0047 | 2.0019 | 2.0044 | -0.0012 |  |
|  | <sup>1</sup>H hyperfine tensors /MHz |  |  |  |  | Assignment |
| A1 | **31.7** | 26.7 | 27.6 | 28.7 | 1.5 | C<sub>β1</sub> |
| A2 | 14.6 | **2.8\*** | 11.9 | 9.8 | -3.5 | H6 |
| A3 | 9.9 | **2.1\*** | 7.5 | 6.5 | -2.2 | H3 |
| A4 | **6.8** | 4.0 | 2.5 | 4.4 | 1.1 | H5 |
| A5 | **4.9** | -2.1 | -2.1 | 0.2 | 2.3 | Dipolar |
| A6 | **3.8** | -1.4 | -1.4 | 0.3 | 1.7 | Dipolar |
| A7 | -1.1 | -1.1 | **4.1** | 0.6 | 1.7 | Dipolar |

\* A<sub>y</sub> component for A2 and A3 is not well defined

**Table S4.** Solid state (frozen solution) EPR data for the ring protons of all tyrosyl radicals characterized in RNR proteins(*68*)

|  |  | <sup>1</sup>H hyperfine tensors /MHz |  |  |  |  |  |
| --- | --- | --- | --- | --- | --- | --- | --- |
|  |  | x | Y | z | iso | aniso | angle |
| E. coli RNR | H2,H6 | 26.7 | **8.4** | 19.6 | 18.2 | -4.9 | ±25 |
|  | H3,H5 | 5.0 | 7.6 | **2.1** | 4.9 | -1.4 | ±44 |
| E. coli RNR | H2,H6 | 26.8 | **7.8** | 19.6 | 18.1 | -5.1 | ±27 |
|  | H3,H5 | 4.8 | 7.5 | **1.1** | 4.5 | -1.7 | ±10 |
| E. coli RNR | H2,H6 | 26.9 | **7.8** | 19.7 | 18.1 | -5.2 | - |
|  | H3,H5 | 4.8 | 7.6 | **1.4** | 4.6 | -1.6 | - |
| Mouse RNR | H2,H6 | 25.1 | **7.0** | 19.0 | 17.0 | -4.7 | ±25 |
|  | H3,H5 | 5.0 | 7.5 | **1.1** | 4.5 | -1.7 | - |

### Assignment of the radical signal seen in the new R2 protein

As described in the main text and above, the radical seen in the new R2 protein has a spectral width and overall structure significantly different from that of all reported tyrosyl radicals. This observation suggests that the ^1^H couplings of the unknown radical species are different from that of a typical tyrosyl radical. Simulation of the EPR lineshape confirm this is the case. An extended discussion is given below that explains why this signal cannot be assigned to a tyrosyl radical or a tyrosyl radical with an additional alkyl substitution(*27*).

#### Reasons to exclude an unmodified tyrosyl (phenoxyl) radical

There are two observations that rule out assigning the new signal to an unmodified tyrosyl radical: i) the magnitude of the beta proton coupling; and ii) the absence of two large, equivalent ring proton couplings. Both of these properties lead to the unknown radical being significantly narrower than all reported tyrosyl radicals.

i) Magnitude of the beta proton coupling: As the crystal structure of the tyrosine in known e.g. the position (dihedral angle 69-72.5) between the CH_2_ side chain and the ring plane, the position of the two beta protons can be inferred. The range of theta angles for proton 1 and 2 are equal to:

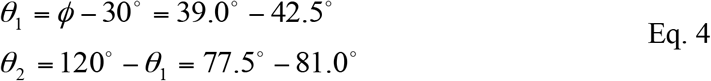

Using Eq. 2 the hyperfine coupling for the beta protons of tyrosine can be obtained. The spin density at C4 for all tyrosines measured falls within 0.34 - 0.43 with the spread correlating with the presence/absence of an H-bond to the phenolic oxygen (*68*). Solution data for para substituted phenoxyl radicals also fall in this range (~0.4).

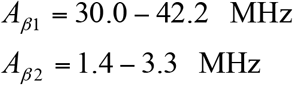

The largest observed coupling (A1_iso_ = 28.5 MHz) is significantly outside this range. The allowed spin density at C4 based on structural constraints is marginally lower 0.29-0.32.

ii) Absence of two large, equivalent ring proton couplings: For all tyrosine radicals reported, two large, symmetry related couplings of the same magnitude are observed (16-18.6 MHz). This has also seen for all phenoxyl radicals in solution, including phenoxyl radicals with additional alkyl substituents. (Strictly speaking the alkyl substitution removes a ring coupling but introduces additional couplings of the same magnitude from the substituent itself. This means that the spin densities within the conjugated ring are approximately the same for all systems and are not strongly affected by alkyl addition.

The two ring coupling observed here fall outside this range (A2 = 9.8, A3 = 6.5 MHz), further demonstrating that the observed radical is not a tyrosine or a tyrosine with an alkyl substitution (in the ortho or meta position) (*27, 68*).

**Figure S9.**
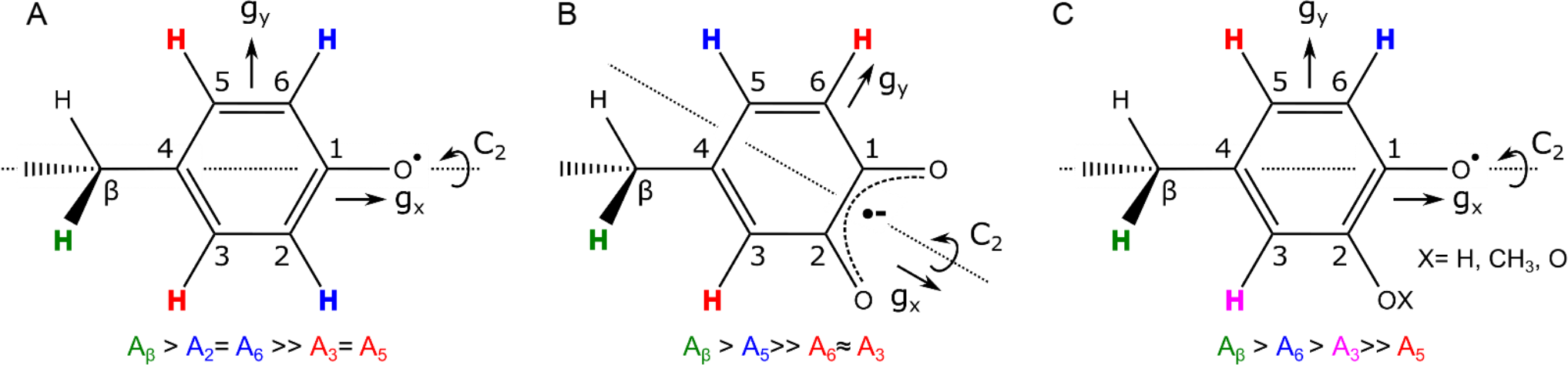
Candidates for the *Mf*R2 radical species.

**Table S5.** Solution EPR data for some p-substituted phenoxyl radicals and o-semiquinone radicals, complied from Stone (1964) and Pillar (1970) (*27. 29*).

| Substituent |  |  | Coupling Constant /MHz |  |  |
| --- | --- | --- | --- | --- | --- |
| 2 | 4 | 6 | a(2,6) | a(4) | a(3,5) |
| H | H | H | 18.6 | 28.3 | 5.0 |
| H | Me | H | 16.8 | (33.5) | 4.1 |
| H | Et | H | 16.8 | (28.4) | 4.2 |
| H | n-Pr | H | 17.1 | (24.4) | 4.1 |
| H | CH<sub>2</sub>Ph | H | 17.1 | (24.4) | 4.1 |
| H | i-Pr | H | 16.8 | (16.8) | ≤3.9 |
| H | s-Bu | H | 16.8 | (12.0) | ≤3.9 |
| H | OMe | H | 16.0 | (4.8) | 2.2 |
| Me | H | Me | (18.2) | 28.3 | 4.6 |
| Et | H | Et | (16.0) | 26.6 | 4.6 |
| Me | Me | H | 18.2 (18.2) | (31.4) | 4.8 |
| t-Bu | Me | H | 18.5 | (31.9) | 4.8 |
| t-Bu | OMe | H | 17.4 | (4.8) | 2.5 |
| OMe | H | H | 11.8 | 23.2 | 5.0, ? |
| OMe | H | OMe | - | 23.2 | 4.2, ? |
| OMe | CHO | H | 12.0 | - | 6.2 |
| O | H | H | -, 2.7 | 10.2 | 2.7, 10.2 |
| O | Me<sup>1</sup> | H | 3.9 | 12.3 | 2.0, 10.2 |
| O | Me<sup>2</sup> | H | 2.8 | 14.3 | 0.7, 11.2 |

<sup>1,2</sup>correspond to different solvent mixtures DMF:water (9:1) and aqueous basic solution

#### The measured proton couplings also rule out its assignment to a genuine semiquinone

An *o*-benzosemiquinone is structurally similar to a tyrosine with an oxygen substituent at C2. For a genuine *o*-benzosemiquinone both oxygens would have approximately the same radical character. and as such the pseudo C2 axis symmetry of the molecule shifts to bisecting the C1-C2, and C4 and C5 bonds. They do however have an overall a lower symmetry to that of the tyrosine upon substitution at the C4 position (i.e. protons of C3 and C6 similar, but not equivalent).

The addition of a second oxygen as part of the conjugated system acts to remove unpaired spin density from the ring, reducing the observed ^1^H hyperfine couplings as is seen for the *Mf*R2 radical. This effect is dramatic, with the C4 spin density dropping to 0.15-0.18(*29*). This value is well outside the range for the spin density at C4 inferred from experiment i.e. 0.29-0.32.

In addition, *o*-benzosemiquinone gives rise to smaller coupling. The largest ring coupling (10 MHz) is now at the at the C5 position i.e. not derived from the protons on the carbons adjacent to those carrying the oxygens. The remaining ring protons typically have coupling half this value i. e. below 4 MHz. While the C5 coupling matches the large ring coupling measured, the absence of a second large coupling disfavours this assignment.

#### A phenoxyl radical with an oxygen substituent at position 2 is the most likely candidate

The observed radical is therefore something in between these two canonical radical types. In this respect literature results on phenoxyl radicals with a methoxy substitution at C2 serve as a guide (*27*). As with benzosemiquinone, the additional oxygen substituent decreases the unpaired spin density shared across the carbon atoms of the ring, leading to lower proton hyperfine coupling - but the magnitude of this effect is far less than seen for an *o*-benzosemiquinone. The spin density at the C4 position for the 2-methoxyphenoxyl as compared to the unsubstituted form, as inferred by the C4 proton coupling) decreases by approximately 20%. Applying this correction to the range seen for tyrosines (0.34-0.43), a rough estimate for the range of spin densities C4 can taken in a methoxy substituted phenoxyl radical is 0.28 to 0.35 (*27*), matching the inferred range for the new radical signal observed.

A radical of this type would also yield two large, non-equivalent ring coupling; as the substitution breaks the C2 axis symmetry of the ring plane, about the C1-C4 axis. As such, the C3 and C6 proton couplings will not be identical. As with a normal phenoxyl, the carbon adjacent to the carbon carrying the phenoxyl radical (C6) has the largest coupling. In the case of a methoxy substitution its only 63% that of the unsubstituted form (18 vs. 11 MHz). The C3, which is instead adjacent to the methoxy group (which has some radical character) is smaller, 27% of its unsubstituted form (18 vs. 5.0 MHz). C5 has only a small coupling. These ^1^H hyperfine couplings (11 and 5 MHz) broadly match those measured for the *Mf*R2 radical (9.8 and 6.5 MHz) (*27*). It is noted though that if the substitute really is a methoxy it introduces three additional, ^1^H couplings of the order of 5 MHz. This additional set of couplings are not observed for the *Mf*R2 radical species, suggesting instead that the substitute is the simpler OH unit. The precise nature of the additional oxygen substituent cannot be resolved solely based on the EPR/ENDOR data. However, it must: i) reduce the unpaired spin density of the ring; ii) break the symmetry of the ring protons; iii) have limited, but not negligible radical character.

**Figure S10.**
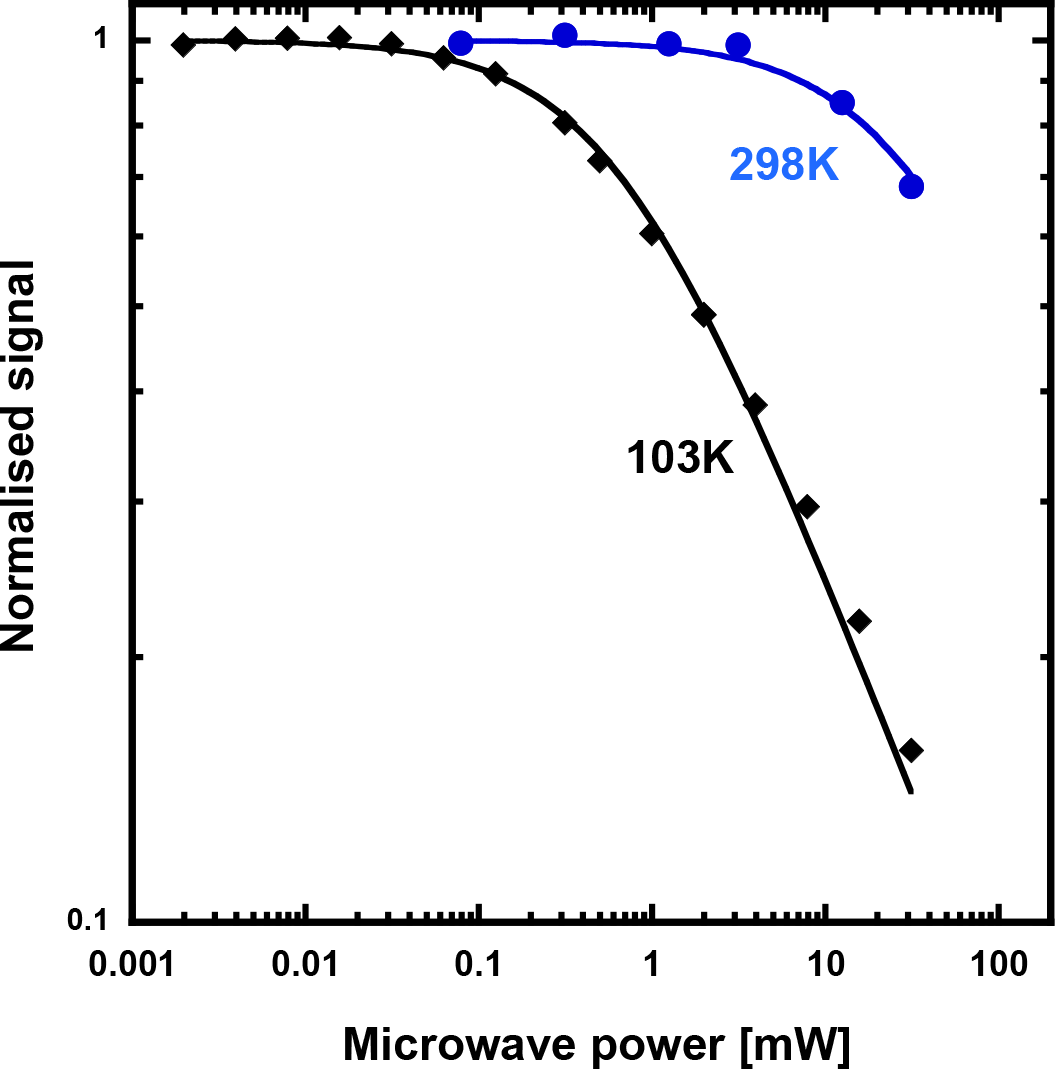
EPR saturation behavior of the *Mf*R2 radical at 103 and 298 K. Saturation curves at different temperatures determine the microwave power at half saturation P_1/2_. The temperature dependence of P_1/2_ gives information about possible relaxing transition metals in the vicinity of the radical. A fast relaxing metal site will give a higher P_1/2_ than for an isolated radical. The microwave saturation behavior of the *Mf*R2 is similar to that for an hv-irradiated tyrosine solution(*30*). Here we evaluate P_1/2_ ≈ 0.6 mW at 103 K and P_1/2_ ≈ 30 mW at 298 K for *Mf*R2. This can be compared to hv Tyr• with P_1/2_ ≈ 0.4mW at 93 K and *E. coli* Tyr• with P_1/2_ ≈ 150 mW at 106 K and not possible to saturate at 298K.

**Table S6.**
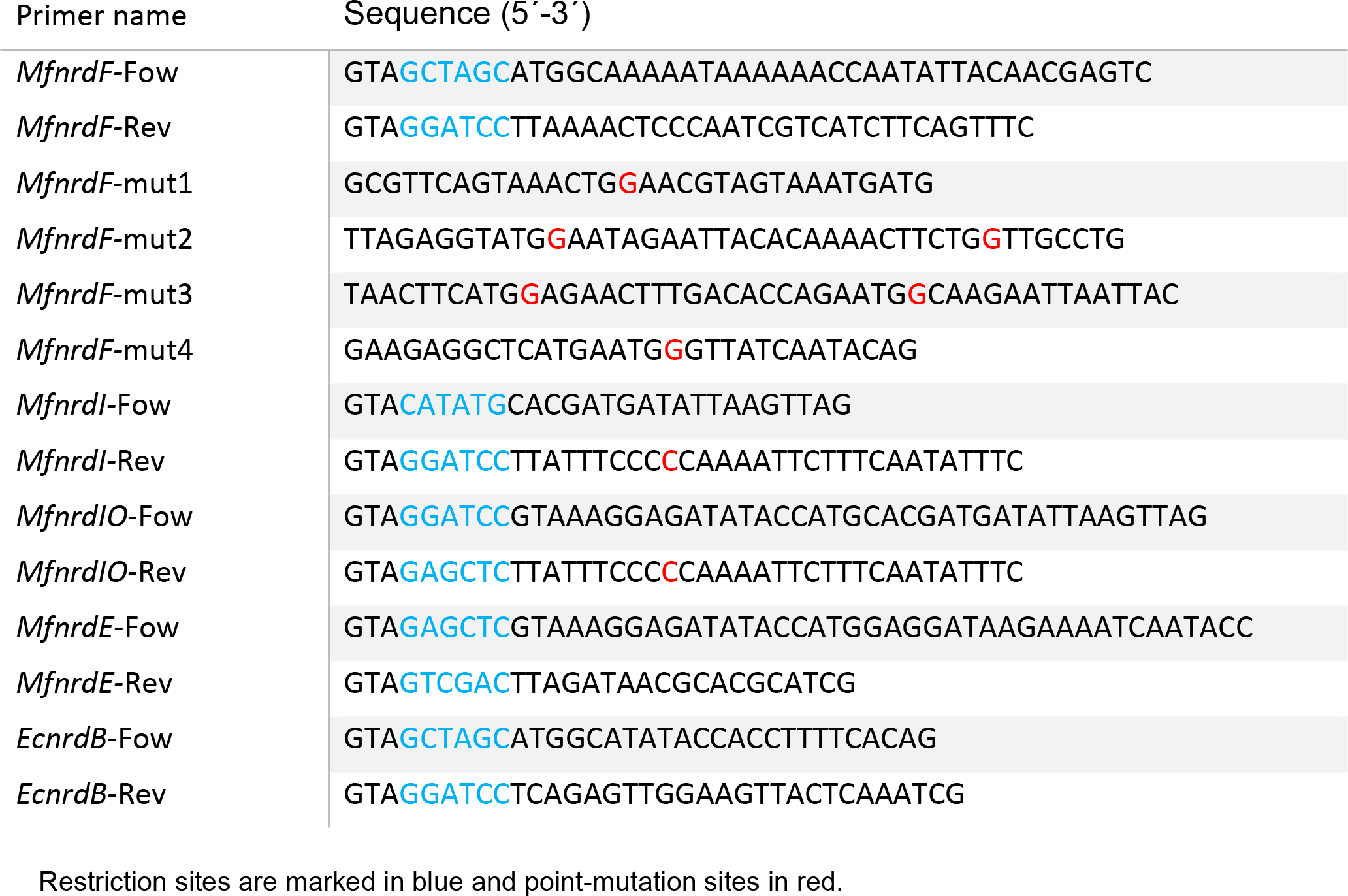
Primers used in this study.

**Figure S11:**
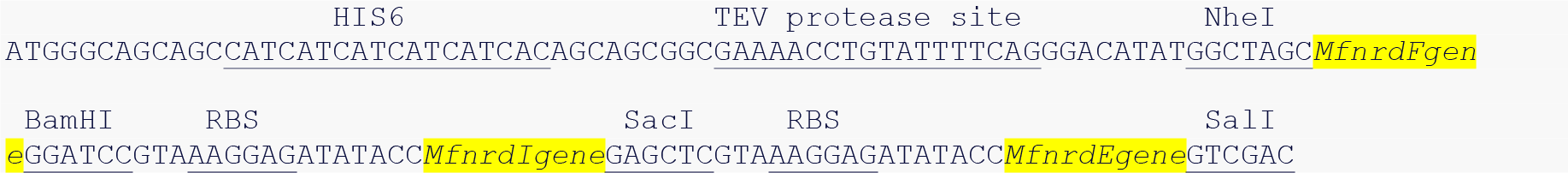
*Mesoplasma florum* class Ie RNR operon construction.

## References

1. A. Hofer, M. Crona, D. T. Logan, B.-M. Sjöberg, DNA building blocks: keeping control of manufacture. Crit. Rev. Biochem. Mol. Biol. 47, 50–63 (2012).

2. P. Nordlund, P. Reichard, Ribonucleotide reductases. Annu. Rev. Biochem. 75, 681–706 (2006).

3. M. B. Mannargudi, S. Deb, Clinical pharmacology and clinical trials of ribonucleotide reductase inhibitors: is it a viable cancer therapy? J. Cancer Res. Clin. Oncol. 143, 1499–1529 (2017).

4. Y. Aye, M. Li, M. J. C. Long, R. S. Weiss, Ribonucleotide reductase and cancer: biological mechanisms and targeted therapies. Oncogene. 34, 2011–2021 (2015).

5. N. C. Brown, R. Eliasson, P. Reichard, L. Thelander, Nonheme iron as a cofactor in ribonucleotide reductase from E. coli. Biochem. Biophys. Res. Commun. 30, 522–527 (1968).

6. D. Lundin, G. Berggren, D. T. Logan, B.-M. Sjöberg, The origin and evolution of ribonucleotide reduction. Life (Basel). 5, 604–636 (2015).

7. M. Huang, M. J. Parker, J. Stubbe, Choosing the right metal: case studies of class I ribonucleotide reductases. J. Biol. Chem. 289, 28104–28111 (2014).

8. P. Nordlund, B. M. Sjöberg, H. Eklund, Three-dimensional structure of the free radical protein of ribonucleotide reductase. Nature. 345, 593–598 (1990).

9. U. Uhlin, H. Eklund, Structure of ribonucleotide reductase protein R1. Nature. 370, 533–539 (1994).

10. J. Stubbe, D. G. Nocera, C. S. Yee, M. C. Y. Chang, Radical initiation in the class I ribonucleotide reductase: Long-range proton-coupled electron transfer? Chem. Rev. 103, 2167–2201 (2003).

11. J. A. Cotruvo, J. Stubbe, Class I ribonucleotide reductases: metallocofactor assembly and repair in vitro and in vivo. Annu. Rev. Biochem. 80, 733–767 (2011).

12. M. Hogbom, Metal use in ribonucleotide reductase R2, di-iron, di-manganese and heterodinuclear--an intricate bioinorganic workaround to use different metals for the same reaction. Metallomics. 3, 110–120 (2011).

13. A. B. Tomter et al., Spectroscopic studies of the iron and manganese reconstituted tyrosyl radical in Bacillus cereus ribonucleotide reductase R2 protein. PLoS ONE. 7, e33436 (2012).

14. D. J. Hirsh, W. F. Beck, G. W. Brudvig, J. B. Lynch, L. Que, Using Saturation-Recovery EPR To Measure Exchange Couplings in Proteins: Application to Ribonucleotide Reductase. Journal of the American Chemical Society. 114, 7475–7481 (1992).

15. M. Hogbom et al., The radical site in chlamydial ribonucleotide reductase defines a new R2 subclass. Science. 305, 245–248 (2004).

16. W. Jiang et al., A manganese(IV)/iron(III) cofactor in Chlamydia trachomatis ribonucleotide reductase. Science. 316, 1188–1191 (2007).

17. G. Berggren, D. Lundin, B.-M. Sjöberg, in Encyclopedia of Inorganic and Bioinorganic Chemistry (American Cancer Society, 2017), pp. 1–17.

18. I. Rozman Grinberg et al., Novel ATP-cone-driven allosteric regulation of ribonucleotide reductase via the radical-generating subunit. Elife. 7, 389 (2018).

19. H. Rose et al., Structural Basis for Superoxide Activation of Flavobacterium johnsoniae Class I Ribonucleotide Reductase and for Radical Initiation by its Dimanganese Cofactor. Biochemistry, acs.biochem.8b00247 (2018).

20. J. A. Cotruvo, T. A. Stich, R. D. Britt, J. Stubbe, Mechanism of assembly of the dimanganese-tyrosyl radical cofactor of class Ib ribonucleotide reductase: enzymatic generation of superoxide is required for tyrosine oxidation via a Mn(III)Mn(IV) intermediate. J. Am. Chem. Soc. 135, 4027–4039 (2013).

21. A. K. Boal, J. A. J. Cotruvo, J. Stubbe, A. C. Rosenzweig, Structural basis for activation of class Ib ribonucleotide reductase. Science. 329, 1526–1530 (2010).

22. G. Berggren, N. Duraffourg, M. Sahlin, B.-M. Sjöberg, Semiquinone-induced maturation of Bacillus anthracis ribonucleotide reductase by a superoxide intermediate. J. Biol. Chem. 289, 31940–31949 (2014).

23. J. E. Martin, J. A. Imlay, The alternative aerobic ribonucleotide reductase of Escherichia coli, NrdEF, is a manganese-dependent enzyme that enables cell replication during periods of iron starvation. Mol. Microbiol. 80, 319–334 (2011).

24. I. Roca, E. Torrents, M. Sahlin, I. Gibert, B.-M. Sjöberg, NrdI essentiality for class Ib ribonucleotide reduction in Streptococcus pyogenes. J. Bacteriol. 190, 4849–4858 (2008).

25. M. Hammerstad, H.-P. Hersleth, A. B. Tomter, A. K. Røhr, K. K. Andersson, Crystal structure of Bacillus cereus class Ib ribonucleotide reductase di-iron NrdF in complex with NrdI. ACS Chem. Biol. 9, 526–537 (2014).

26. E. J. Land, G. Porter, Primary photochemical processes in aromatic molecules. Part7.—Spectra and kinetics of some phenoxyl derivatives. Trans. Faraday Soc. 59, 2016–2026 (1963).

27. T. J. Stone, W. A. Waters, Aryloxy-radicals. Part 1. Electron spin resonance spectra of radicals from some substituted monojiydric phenols. Journal of the Chemical Society (Resumed), 213–218 (1964).

28. W. G. B. Huysmans, W. A. Waters, Aryloxy-radicals. Part VI. Measurement of the electron spin resonance spectra of short-lived substituted phenoxy-radicals in benzene solution. Journal of the Chemical Society B: Physical Organic. 0, 1047–1049 (1966).

29. J. Pilar, Electron paramagnetic resonance study of 4-Alkyl-o-benzosemiquinones. Journal of Physical Chemistry. 74, 4029–4037 (1970).

30. M. Sahlin et al., Magnetic interaction between the tyrosyl free radical and the antiferromagnetically coupled iron center in ribonucleotide reductase. Biochemistry. 26, 5541–5548 (1987).

31. A. Ehrenberg, P. Reichard, Electron spin resonance of the iron-containing protein B2 from ribonucleotide reductase. J. Biol. Chem. 247, 3485–3488 (1972).

32. L. Thelander, B. Larsson, J. Hobbs, F. Eckstein, Active site of ribonucleoside diphosphate reductase from Escherichia coli. Inactivation of the enzyme by 2’-substituted ribonucleoside diphosphates. J. Biol. Chem. 251, 1398–1405 (1976).

33. R. Eliasson, E. Pontis, F. Eckstein, P. Reichard, Interactions of 2“-modified azido-and haloanalogs of deoxycytidine 5-”triphosphate with the anaerobic ribonucleotide reductase of Escherichia coli. J. Biol. Chem. 269, 26116–26120 (1994).

34. B. M. Sjöberg, A. Gräslund, F. Eckstein, A substrate radical intermediate in the reaction between ribonucleotide reductase from Escherichia coli and 2“-azido-2-”deoxynucleoside diphosphates. J. Biol. Chem. 258, 8060–8067 (1983).

35. M. R. Seyedsayamdost, J. Stubbe, Site-specific replacement of Y356 with 3,4-dihydroxyphenylalanine in the beta2 subunit of E. coli ribonucleotide reductase. J. Am. Chem. Soc. 128, 2522–2523 (2006).

36. H. S. Raper, Note on the oxidation of tyrosine, tyramine and phenylalanine with hydrogen peroxide. Biochem. J. 26, 2000–2004 (1932).

37. G. Cohen, S. Yakushin, D. Dembiec-Cohen, Protein L-dopa as an index of hydroxyl radical attack on protein tyrosine. Anal. Biochem. 263, 232–239 (1998).

38. J. Quintero-Saumeth, D. A. Rincón, M. Doerr, M. C. Daza, Concerted double proton-transfer electron-transfer between catechol and superoxide radical anion. Phys Chem Chem Phys. 19, 26179–26190 (2017).

39. B. M. Sjöberg, P. Reichard, A. Gräslund, A. Ehrenberg, The tyrosine free radical in ribonucleotide reductase fromEscherichia coli. J. Biol. Chem. 253, 6863–6865 (1978).

40. M. I. Hood, E. P. Skaar, Nutritional immunity: transition metals at the pathogen-host interface. Nat. Rev. Microbiol. 10, 525–537 (2012).

41. S. R. Eddy, Accelerated Profile HMM Searches. PLoS Comput. Biol. 7, e1002195 (2011).

42. R. C. Edgar, Search and clustering orders of magnitude faster than BLAST. Bioinformatics. 26, 2460–2461 (2010).

43. C. B. Do, M. S. P. Mahabhashyam, M. Brudno, S. Batzoglou, ProbCons: Probabilistic consistency-based multiple sequence alignment. Genome Res. 15, 330–340 (2005).

44. A. Criscuolo, S. Gribaldo, BMGE (Block Mapping and Gathering with Entropy): a new software for selection of phylogenetic informative regions from multiple sequence alignments. BMC Evol. Biol. 10, 210 (2010).

45. A. Stamatakis, RAxML version 8: a tool for phylogenetic analysis and post-analysis of large phylogenies. Bioinformatics. 30, 1312–1313 (2014).

46. C. Loderer et al., A unique cysteine-rich zinc finger domain present in a majority of class II ribonucleotide reductases mediates catalytic turnover. J. Biol. Chem. 292, 19044–19054 (2017).

47. W. Kabsch, XDS. Acta Crystallogr. Sect. D-Biol. Crystallogr. 66, 125–132 (2010).

48. A. J. McCoy et al., Phaser crystallographic software. J. Appl. Crystallogr. 40, 658–674 (2007).

49. M. E. Andersson et al., Structural and mutational studies of the carboxylate cluster in iron-free ribonucleotide reductase R2. Biochemistry. 43, 7966–7972 (2004).

50. P. D. Adams et al., PHENIX: a comprehensive Python-based system for macromolecular structure solution. Acta Crystallogr D Biol Crystallogr. 66, 213–221 (2010).

51. P. Emsley, K. Cowtan, Coot: model-building tools for molecular graphics. Acta Crystallogr D Biol Crystallogr. 60, 2126–2132 (2004).

52. V. B. Chen et al., MolProbity: all-atom structure validation for macromolecular crystallography. Acta Crystallogr. Sect. D-Biol. Crystallogr. 66, 12–21 (2010).

53. E. Krissinel, K. Henrick, Secondary-structure matching (SSM), a new tool for fast protein structure alignment in three dimensions. Acta Crystallogr. Sect. D-Biol. Crystallogr. 60, 2256–2268 (2004).

54. E. Reijerse, F. Lendzian, R. Isaacson, W. Lubitz, A tunable general purpose Q-band resonator for CW and pulse EPR/ENDOR experiments with large sample access and optical excitation. Journal of Magnetic Resonance. 214, 237–243 (2012).

55. D. Franke et al., ATSAS 2.8: a comprehensive data analysis suite for small-angle scattering from macromolecular solutions. J Appl Crystallogr. 50, 1212–1225 (2017).

56. P. V. Konarev, V. V. Volkov, A. V. Sokolova, M. H. J. Koch, D. I. Svergun, PRIMUS: A Windows PC-based system for small-angle scattering data analysis. J Appl Crystallogr. 36, 1277–1282 (2003).

57. D. Franke, D. I. Svergun, DAMMIF, a program for rapid ab-initio shape determination in small-angle scattering. J Appl Crystallogr. 42, 342–346 (2009).

58. V. V. Volkov, D. I. Svergun, Uniqueness of ab initio shape determination in small-angle scattering. J. Appl. Cryst. (2003). 36, 860-864, vol. 36, pp. 860–864.

59. D. Svergun, C. Barberato, M. H. Koch, CRYSOL - A program to evaluate X-ray solution scattering of biological macromolecules from atomic coordinates. J Appl Crystallogr. 28, 768–773 (1995).

60. M. Sielaff et al., Evaluation of FASP, SP3, and iST Protocols for Proteomic Sample Preparation in the Low Microgram Range. J. Proteome Res. 16, 4060–4072 (2017).

61. C. S. Hughes et al., Ultrasensitive proteome analysis using paramagnetic bead technology. Mol. Syst. Biol. 10, 757–757 (2014).

62. J. K. Eng, A. L. McCormack, J. R. Yates, An approach to correlate tandem mass spectral data of peptides with amino acid sequences in a protein database. J. Am. Soc. Mass Spectrom. 5, 976–989 (1994).

63. S. Na, N. Bandeira, E. Paek, Fast multi-blind modification search through tandem mass spectrometry. Mol. Cell Proteomics. 11, M111.010199 (2012).

64. M. C. Chambers et al., A cross-platform toolkit for mass spectrometry and proteomics. Nat. Biotechnol. 30, 918–920 (2012).

65. A. Carrington, A. D. McLachlan, Introduction to magnetic resonance with applications to chemistry and chemical physics (1967).

66. S. Stoll, A. Schweiger, EasySpin, a comprehensive software package for spectral simulation and analysis in EPR. J. Magn. Reson. 178, 42–55 (2006).

67. S. Stoll, R. D. Britt, General and efficient simulation of pulse EPR spectra. Phys Chem Chem Phys. 11, 6614–6625 (2009).

68. D. A. Svistunenko, C. E. Cooper, A new method of identifying the site of tyrosyl radicals in proteins. Biophys. J. 87, 582–595 (2004).

